# Improving the fidelity of uridine analog incorporation during *in vitro* transcription

**DOI:** 10.1101/2022.04.12.488100

**Authors:** Tien-Hao Chen, Vladimir Potapov, Nan Dai, Jennifer L. Ong, Bijoyita Roy

## Abstract

*In vitro* transcribed synthetic messenger RNAs (mRNAs) represent a novel therapeutic modality and are currently being evaluated for a wide range of clinical indications. To overcome the inherent immunogenicity of the synthetic mRNAs, as well as to increase the therapeutic efficacy of the molecules, RNA sequence optimization is routinely performed and modified uridine analogs—such as pseudouridine (Ψ) and *N*^1^-methyl-pseudouridine (m1Ψ), are incorporated in the synthetic mRNA. To decipher the fidelity with which these modifications are incorporated during the *in vitro* transcription (IVT) process, here, we compared, the incorporation fidelity of uridine, Ψ, or m1Ψ in multiple RNA sequences with different single-subunit DNA-dependent RNA polymerases (ssRNAPs). By comparing the incorporation of each modified base to that of the unmodified equivalent, we demonstrate that m1Ψ is incorporated with higher fidelity than Ψ. Furthermore, the various ssRNAPs exhibit different error rates; however, the spectrum of mutations observed between the RNAPs is similar. We also show that the array of nucleotide misincorporation is not dependent on the template DNA sequence context and that the distribution of these misincorporated nucleotides is not localized to any specific region along the length of the RNA. Based on our findings, we introduce a novel protocol to improve uridine analog incorporation—without affecting total RNA yield—during IVT. Our proof-of-concept experiments and protocol for higher-fidelity incorporation of uridine analogs during IVT provide guidelines when choosing ssRNAPs for the generation of modified uridine—containing mRNAs *in vitro*.

## Introduction

Synthetic messenger RNAs (mRNAs) are a novel modality for vaccines, and they are also being evaluated as a vector for therapeutics [1–3]. Despite there being several advantages over conventional protein-based approaches, mRNA-based therapeutics are still in early stages of development. Instability of the synthetic mRNAs and the immune responses generated against these synthetic molecules have been key hurdles in the adaptation of this technology, particularly for therapeutic applications where prolonged expression from the synthetic molecule is desirable and repeated dosing of the drug product is required [1, 2]. The use of chemically modified bases in synthetic mRNAs is an innovation that has allowed for both an ameliorated immune response to the synthetic molecules and increased protein expression from the mRNA, thereby providing an unprecedented opportunity to use synthetic mRNAs as a novel class of therapeutics for a wide range of indications, including two approved vaccines against COVID-19 [4, 5]. It has been shown that the incorporation of pseudouridine (ψ), N1-methyl-pseudouridine (m1ψ), 5-methylcytosine (m5C), N6-methyladenosine (m6A) and 2-thiouridine (s2U) into synthetic mRNAs results in reduced immune responses and increased protein expression *in vivo* [6–14]. Pseudouridine-modified mRNAs have been shown to result in reduced activation of 2’-5’-oligoadenylate synthetase (OAS), RNA-dependent protein kinase (PKR), and toll-like receptors [10]. Investigation of ψ derivatives with improved pharmacological properties led to the identification of m1ψ, the current benchmark for synthetic mRNA-based applications. m1ψ is a naturally occurring modification found in 18S rRNA and tRNAs [15–17], and similar to ψ-modified synthetic mRNAs, the presence of m1ψ in synthetic mRNAs has been demonstrated to result in reduced activation of RNA sensors in cells [8, 12, 14, 18]. Furthermore, the presence of m1ψ in synthetic mRNAs show increased translation efficiency in cell-free extracts, multiple mammalian cell lines, and mouse models [8, 9, 12–14, 18–20]. The exact mechanism of how m1ψ enhances translation is not well understood but it has been demonstrated that presence of m1ψ, alone or in combination with other chemical modifications, can alter ribosome transit time on the modified mRNA, and can increase the mRNA half-life by altering the secondary structure of the synthetic mRNA [7, 14, 21, 22].

Current methods to incorporate modified nucleotides in synthetic mRNAs include complete substitution of the standard nucleotide with a chemically modified nucleotide during the process of *in vitro* transcription (IVT) by single-subunit DNA-dependent RNA polymerases (ssRNAPs; such as T7, T3, and SP6 RNAP) [1]. In contrast to endogenous mRNAs, in which modified nucleotides occur in specific positions in the mRNA [23, 24], in synthetic mRNAs, the modified nucleotide is present at almost every position where the naturally occurring nucleobase would be. This complete substitution approach is preferred from a regulatory perspective because it results in less molecule-to-molecule variation in the positions of the modified nucleotides along the synthetic mRNA. The exact implications of incorporating the chemical modifications throughout the body of the mRNA is still under investigation. For attaining the maximum effectiveness from the drug product and to ensure that the expression from the synthetic molecule is optimal, it is critical that the modified nucleotide is incorporated in the right place, be compatible with the functional elements of the mRNA, and does not alter the biological function of the synthetic mRNA. Numerous studies have demonstrated that ssRNAPs can incorporate chemically modified nucleotides into RNA [7, 10, 25], but it is unclear whether all ss-RNAPs incorporate the modified nucleotides with comparable fidelity.

We previously established a Pacific Biosciences Single Molecule Real-Time (SMRT) sequencing-based assay to determine the combined transcription and reverse transcription errors and the effects of RNA modifications [26]. In the previous study, T7 RNAP exhibited higher combined error rates in the synthesis of ψ-, m6A- and 5-hydroxymethylcytidine (hm5C)-modified RNAs, with increased misincorporation of ψ across from dT templated bases. In light of the widespread use of m1ψ for mRNA vaccines, in the current study, we focused on understanding the fidelity of incorporation of m1ψ during *in vitro* transcription using multiple ssRNAPs. The fidelity with which m1ψ is incorporated during *in vitro* transcription is a timely question to address because both the SpikeVax (mRNA-1273) and Comirnaty (BNT162b2) SARS-CoV2 mRNA vaccines are synthesized by substituting uridine with m1ψ throughout the body of the mRNA and both have demonstrated efficacy greater than 94% [4, 5]. We investigated the fidelity of m1ψ incorporation with three commonly used ssRNAPs (T7, T3 and SP6 RNAPs) and compared it to the fidelity with which uridine and ψ are incorporated. Using four different RNA sequences, including two functional mRNA sequences, we analyzed the effects of sequence context in promoting misincorporation of uridine analogs and identified rA→rU substitution errors as the predominant error when uridine analogs (m1ψ and ψ) are present in the reaction. Finally, based on the nature of the substitution errors observed, we combine sequence optimization of the synthetic mRNA together with an altered RNA synthesis process to reduce the uridine analog incorporation error during *in vitro* transcription. These proof-of-concept experiments establish a novel method to synthesize mRNAs with improved fidelity of uridine analog incorporation and provide considerations for choosing ssRNAPs for the generation of modified nucleotide-containing mRNAs *in vitro*.

## Materials and Methods

### Oligonucleotides and DNA template sequences used in this study

All of the oligonucleotides for *in vitro* transcription and reverse transcription were synthesized by Integrated DNA Technologies (IDT, Coralville IA) and the sequences are available in **Table S19**. The DNA template sequences for *in vitro* transcription are also available in **Table S20**.

### Generation of DNA templates for *in vitro* transcription (IVT) of long RNAs

Generation of the DNA templates for the artificial RNA sequences were described earlier [26]. For *in vitro* transcription reactions with SP6 or T3 RNA polymerase, the corresponding promoter sequences were inserted in the DNA templates using Q5 Site-Directed Mutagenesis Kit (E0554, New England Biolabs). DNA templates encoding for functional mRNAs, RNA2 (*Cypridina* luciferase mRNA; 1707 nucleotides) and RNA6 (part of BNT162b/Comirnaty mRNA; 4187 nucleotides) [27, 28], were synthesized by GenScript Inc. (GenScript, Piscataway NJ) and introduced into standard high-copy plasmids. DNA templates for uridine-depleted artificial sequences (RNA7, RNA8, RNA9) were also synthesized by GenScript and introduced into standard high-copy plasmids. The plasmids were propagated in *E. coli* (C2987, New England Biolabs) and purified with the Monarch Plasmid Miniprep Kit (T1010, New England Biolabs). Plasmids were digested with restriction enzymes to generate linearized templates for *in vitro* transcription. The linearized plasmids were treated with PreCR Repair Mix (M0309, New England Biolabs) and purified with the Monarch PCR & DNA Cleanup Kit (T1030, New England Biolabs).

For short RNAs (30 nucleotide RNA3 and 60 nucleotide RNA4), double-stranded DNA templates were generated by annealing DNA oligonucleotides for template and non-template strands in annealing buffer (10 mM Tris pH 7.5, 50 mM NaCl, 1 mM EDTA) by heating at 95°C for 5 min followed by gradual cooling to room temperature.

### *In vitro* transcription (IVT)

*In vitro* transcription reactions were performed according to standard protocols provided with the high-yield *in vitro* transcription kits (E2040 and E2070, New England Biolabs), consisting of 40 mM rNTP (pH buffered with sodium phosphate) for T7 RNA polymerase and 20 mM rNTP (pH buffered with Tris) for SP6 RNA polymerase. For modified RNAs, UTP was replaced with either pseudouridine-5’-triphosphate (N-1019, TriLink Biotechnologies) or N^1^-Methylpseudouridine-5’-Triphosphate (N-1081, TriLink Biotechnologies). For the uridine-depleted sequences (RNA6, RNA7 and RNA8), two sets of reactions were performed—one in which all the rNTPs were present in equimolar ratio and another in which the rNTPs were altered to match the template sequence composition. Following IVT, the DNA template was digested with Turbo DNase (AM2238, Invitrogen) digestion at 37°C for 30 minutes and then purified with the Monarch RNA Cleanup Kit (T2050, New England Biolabs) for long RNA (1020 nucleotides to 4187 nucleotides) or with the Oligo Clean-Up and Concentration Kit (34100, Norgen Biotek Inc) for short RNA (30 nucleotides to 60 nucleotides).

Low-yield IVT reaction conditions were carried out with 70 nM T7 RNA polymerase (M0251, New England Biolabs), 20 mM or 10 mM rNTP, in buffer containing 40 mM Tris-HCl pH 7.9, 20 mM MgCl_2_, 1 mM DTT, 2 mM spermidine supplemented with 1 unit/μL RNase inhibitor, murine (M0314, New England Biolabs) and 2.5 unit/μL inorganic pyrophosphatase (M2403, New England Biolabs) at 37°C for two hours. DNA template removal and cleanup were performed as described above.

### Bioanalyzer for RNA size distribution and integrity

Eluted RNA samples from the *in vitro* transcription reactions were diluted based on concentrations measured on a Nanodrop spectrophotometer (13-400-519, Thermo Fisher Scientific) and denatured at 70°C for 2 minutes and snap-cooled on ice. 250 ng RNA samples were analyzed with RNA 6000 Nano kits (5067, Agilent Technologies) and the integrity and size distribution of the RNA was assessed using mRNA Nano series 2 assay (G2938, Agilent Technologies).

### Nucleoside digestion of RNA and UHPLC-MS analyses to assess modification incorporation

Purified modified RNA and RNA without any chemical modification were digested with the Nucleoside digestion mix (M0649, New England Biolabs) at 37°C for 1 hour. Base composition analysis was performed by Liquid Chromatography-Mass Spectrometry (LC-MS) using an Agilent 1290 Infinity II UHPLC equipped with G7117A Diode Array Detector and 6135XT MS Detector, on a Waters Xselect HSS T3 XP column (2.1 × 100 mm, 2.5 μm) with the gradient mobile phase consisting of methanol and 10 mM ammonium acetate buffer (pH 4.5).

### Intact Mass Spectroscopy analysis for short RNA oligonucleotides

Purified samples of RNA3 and RNA4 with different uridine modifications were analyzed at Novatia, LLC using on-line desalting, flow injection electrospray ionization on an LTQ-XL ion trap mass spectrometer (Thermo Fisher Scientific) and analyzed with ProMass Deconvolution software.

### First and second strand cDNA synthesis for PacBio

The cDNA synthesis was performed as described before with a modified cleanup step using Monarch PCR & DNA Cleanup Kit (T1030, New England Biolabs) [26]. Sequences of forward and reverse oligonucleotides used are provided in the **Table S19**.

### Pacific Biosciences SMRTbell library preparation and sequencing

The library preparation for sequencing on RSII system was performed as described before [26]. For the sequencing on the Sequel platform, about 1.5 μg cDNA was treated with NEBNext End Repair Module (E6050, New England Biolabs) at room temperature for 5 minutes, followed by purification with Monarch PCR & DNA Cleanup Kit (T1030, New England Biolabs). The end-repaired cDNA was ligated with 2 μL barcoded adaptor (100-466-000, Pacific Biosciences) with T4 DNA Ligase (M0202, New England Biolabs) in 50 μL reaction volume at room temperature for 1 hour, followed by purification with Monarch PCR & DNA Cleanup Kit (T1030, New England Biolabs). The un-ligated adaptor and cDNA were digested with *E. coli* Exonuclease III (M0206, New England Biolabs) and Exonuclease VII (M0379, New England Biolabs) in 1X standard Taq buffer at 37°C for 1 hour, followed by cleaning up with Monarch PCR & DNA Cleanup Kit (T1030, New England Biolabs). The ligated DNA was repaired with PreCR Repair Mix (M0309, New England Biolabs) at 37°C for 30 minutes. The libraries were purified with 0.6X volume of AMPure PB beads (100-265-900, Pacific Biosciences) and pooled for sequencing runs. SMRT Link was used to generate the protocol for primer annealing, polymerase binding [Sequel Binding Kit 3.0 (101-613-900, Pacific Biosciences)], cleanup and final loading to three SMRT Cells LR and sequencing using Sequel system.

### Data Analysis

Analysis of sequencing data was performed as previously described [26]. In short, high-accuracy consensus sequences were built for the first and second strand for each sequenced double-stranded DNA. The consensus sequences were aligned to the reference sequence and base substitutions, deletions, and insertions were determined. The first strand error rates were determined by comparing the first strand consensus sequences to the reference sequence (RNA strand), and mutations were required to be present in both strands. The average error and standard deviation were then calculated for each template and enzyme: for RNA1 and RNA5 (1122- and 1124-nucleotide synthetic sequence that includes all possible four-base combinations), one measurement was performed and the error rates were combined and referred as RNA1/RNA5. For other RNA templates, two independent repeats were performed. The relative fold change was calculated for each substitution as (M—U) / U, where M is the substitution rate on modified RNA (m1ψ or ψ) and U is the substitution rate on unmodified RNA. The local context of base substitutions was extracted, and sequence logos were built using WebLogo software [29]. Also, the frequency of mutations at each position was plotted to examine distribution of errors along the reference sequence.

## Results

### T7 and SP6 RNAPs incorporate m1ψ efficiently

The efficiency of m1ψ incorporation during *in vitro* transcription was investigated in four different RNA substrates of varying length and sequence. The base composition of the synthesized RNA was analyzed with ultra-high performance liquid chromatography coupled with mass spectrometry. For the long synthetic RNAs with length ranging from 1122 to 4178 nucleotides, the integrity of the RNA was determined using Bioanalyzer. Synthesis of full-length RNAs of expected sizes were observed in reactions performed in the presence of m1ψ with both T7 RNAP and SP6 RNAP (**Fig. S1A**). Furthermore, similar to ψ, total m1ψTP incorporation was as expected in RNA sequences RNA1 (a 1122 nucleotide synthetic sequence that includes all possible four-base combinations) [30] and RNA2 (Cypridina luciferase mRNA sequence) when *in vitro* transcription was performed with T7 RNAP (**Fig. S1B**). In order to determine that the RNA synthesized in presence of m1ψTP is indeed of expected length, we subjected two short RNAs corresponding to 30 nucleotide (RNA3) and 60 nucleotides (RNA4) to intact mass spectrometry analyses (**Fig. S2**). Irrespective of which uridine analog was present in the reaction, the mass of the predominant species observed in reactions performed with T7 RNAP correspond to the run-off transcripts and few non-templated additions were observed [31] suggesting that similar to ψ and uridine, m1ψ is incorporated efficiently during *in vitro* transcription and the modifications did not disrupt the synthesis of full-length run-off products.

### m1ψ is incorporated with higher fidelity than ψ by T7 RNAP

To determine the fidelity of m1ψ incorporation during *in vitro* transcription, we adapted the Pacific Biosciences Single Molecule Real-Time (SMRT) sequencing-based assay (**Fig. S3**) to the PacBio Sequel I system to enable higher sequencing capacity and multiplexing [26]. The *in vitro* transcribed RNAs were reverse transcribed into double-stranded cDNA using ProtoScript II reverse transcriptase (RT) and sequenced in the SMRT sequencing platform. We analyzed the errors in the first strand that stem from combined RNAP and RT error, referred hereafter as combined errors. We first determined the combined errors in two synthetic sequences (RNA1 and RNA5) that represent all possible four-base combinations (templates described as DNA-1 and DNA-2 previously [26]). The error rates of pooled RNA1 and RNA5 (referred as RNA1/RNA5) in reactions with canonical uridine, using the Sequel I system was observed to be 6.4 ± 0.4 x 10^-5^ error/base as compared to 5.6 ± 0.8 x 10^-5^ error/base using the PacBio RSII system (**Table S1**) [26]. Since the combined error rates were comparable between the two platforms, we used the Sequel I system for the subsequent experiments.

We next tested the error rates in reactions with m1ψ for RNA1 and RNA5 sequences when *in vitro* transcription reactions are performed with T7 RNAP under standard high-yield reaction conditions. Total combined error rates in m1ψ-containing reactions were observed to be 8.0 ± 0.3 x 10^-5^ as compared to 6.4 ± 0.4 x 10^-5^ error/base for uridine-containing reactions and 1.1 ± 0.2 x 10^-4^ errors/base for ψ-containing reactions (**Fig. 1A, Table S2**). The combined error rates in ψ-modified RNAs were observed to be greater than unmodified RNAs, while that in m1ψ-modified RNAs were comparable to unmodified RNAs (**Fig. 1A, Table S2**). To ensure that the observed combined error rates are not dependent on the sequence of the RNA, we also analyzed the combined error rates in two functional mRNAs (RNA2 encoding *Cypridina* luciferase and RNA6 encoding part of BNT162b/Comirnaty mRNA) [27, 28]. Similar to what was observed with the two synthetic sequences, we observed error rates of 6.1 ± 0 x 10^-5^ to 1.4 ± 0.3 x 10^-4^ errors/base for RNA2 (*Cypridina* luciferase) and from 4.7 ± 0.1 x 10^-5^ to 1.3 ± 0 x 10^-4^ errors/base for RNA6 (part of BNT162b/Comirnaty mRNA sequence) (**Fig. 1A, Table S2**). Interestingly, in presence of ψ, error/base was consistently observed to be two-fold higher than that observed with unmodified luciferase or BNT162b/Comirnaty mRNA. In contrast, consistent with RNA1 and RNA 5, when m1ψ was present in the reaction, the error/base was observed to be less than that observed with ψ-modified luciferase or BNT162b/Comirnaty mRNA (**Fig. 1A, Table S2**).

**Figure 1.**
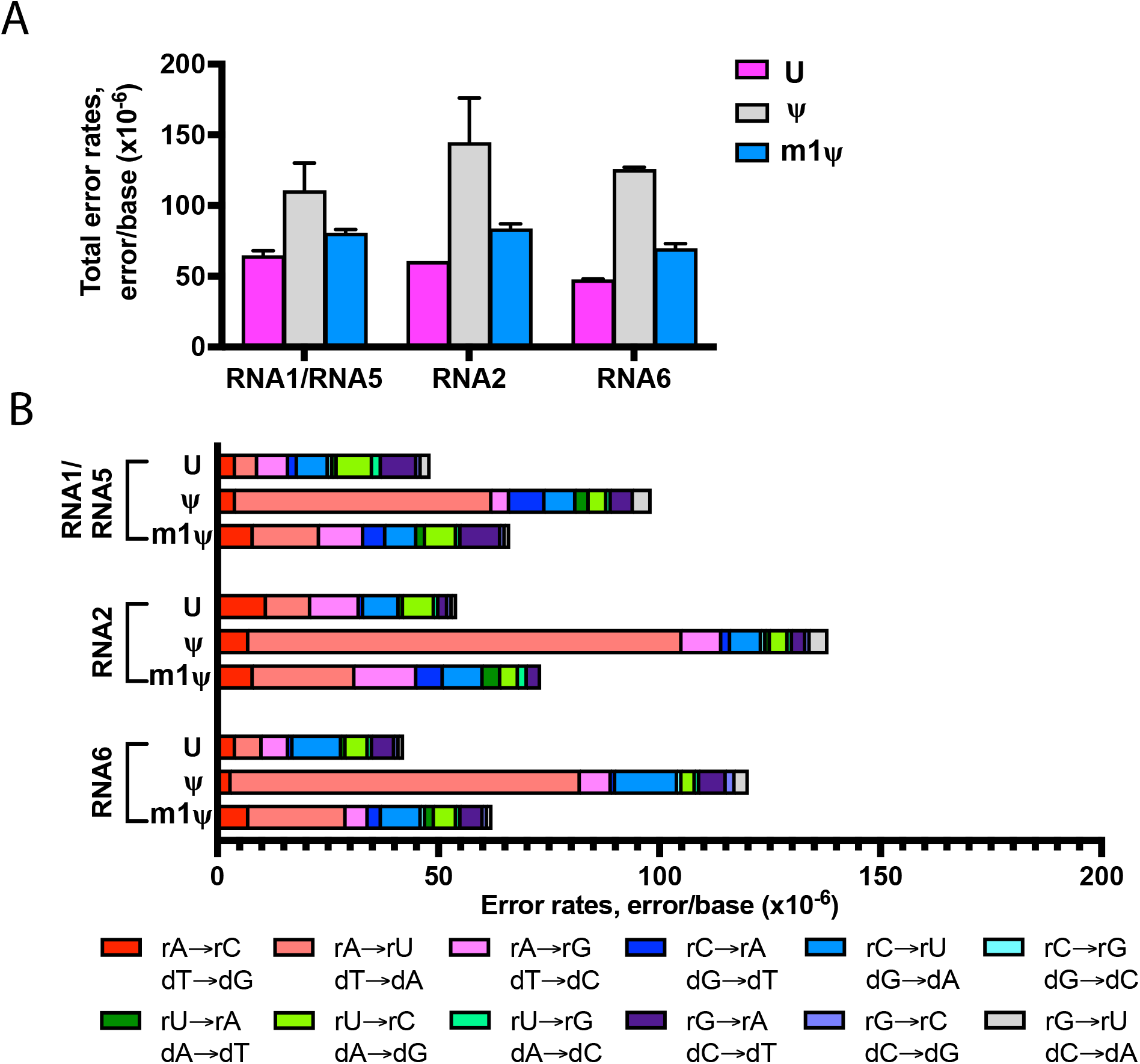
Uridine, ψ, and m1ψ are incorporated by T7 RNA polymerase with varying error rates during *in vitro* transcription. The RNA templates synthesized with T7 RNA polymerase were converted into cDNA with ProtoScript II reverse transcriptase followed by library preparation and sequencing. First-strand errors represented here stem from the combination of RNA polymerase and reverse transcriptase errors (**A**). RNA1 and RNA5 are artificial RNA sequences that have been permutated to include every four-base combination, RNA2 is a functional mRNA encoding Cypridina luciferase and RNA6 contains part of Comirnaty sequence. (**B**) Base substitution error profile observed for unmodified and modified RNA sequences are represented. Colors indicate specific substitution errors observed.

In order to understand whether the nature of the errors that are introduced when the reactions are performed with different uridine analogs is similar, we compared the error profile observed in the four RNA sequences. Irrespective of the sequence of the RNA or the nature of the uridine analog used in the reaction, base substitution was observed to account for the predominant errors ranging from 73% to 96% of total errors (**Table S2**). To further understand if a specific substitution is more prevalent than another and if there are differences when m1ψ is present in the reaction, we analyzed the substitution profile (**Fig. 1B, Table S3**). For unmodified RNA, no distinct substitution errors were observed for any of the sequences (**Fig. 1B, Table S3**). In contrast, a significant increase in rA→rU/dT→dA substitutions were observed when reactions were performed with either m1ψ or ψ (**Fig. 1B, Table S3**), likely due to m1ψTP incorporated in place of rATP (opposite dT) by T7 RNA polymerase as it is less likely that m1ψ would have a global effect on the incorporation of other bases. The rA→rU/dT→dA error rates in ψ-modified RNAs were about nine-to 12-fold higher than unmodified RNAs. The rA→rU/dT→dA substitution was also apparent (upto three-fold greater than unmodified RNAs) in m1ψ-modified RNAs but was observed to be less compared to ψ-modified counterpart (**Fig. 1B, Table S3**).

Even though the nature of the base substitution errors were similar for all four RNA sequences, we further investigated if there is a specific sequence context within these RNA sequences that have a higher propensity of association with the rA→rU/dT→dA substitution errors observed. Neighboring sequence analyses of the rA→rU/dT→dA sites demonstrated a pronounced 5’ rC sequence to be present when the rA→rU/dT→dA substitution was observed in the ψ-modified RNAs (**Fig. S4**). Furthermore, the rA→rU sites were observed to be distributed throughout the RNA and specific hot spots for the substitutions were not observed for any of the RNA sequences, suggesting that the errors observed do not have a sequence context dependency (**Fig. S5**).

### SP6 RNAP incorporates m1ψ with higher fidelity than ψ; overall error rates in reactions performed with SP6 RNAPs are higher than those performed with T7 RNAP

We next sought to compare different RNAPs to determine if m1ψ is incorporated with varied fidelity by different ssRNAPs and if the differences observed in error rates in ψ- and m1ψ-incorporating RNAs synthesized with T7 RNAP are also observed with other ssRNAPs, T3 and SP6 RNAPs. SP6 RNAP shares 32% identity to T7 RNAP and is also used for generating synthetic mRNAs for therapeutic applications [25]. On the other hand, T3 RNAP is 82% identical to T7 RNAP. The total combined error rates observed in reactions performed with T3 RNAP were comparable to reactions performed with T7 RNAP suggesting that these two closely related RNAPs have similar fidelity profile (**Table S4)**. On the other hand, comparison of the total combined error rates observed from modified and unmodified RNAs synthesized with SP6 RNAP demonstrated two-fold to three-fold higher error rates than those observed with T7 RNAP (**Figs. 1A and 2A, Tables S2, S4 and S5**). For unmodified RNA1/RNA5, the total combined error rates observed when reactions were performed with SP6 RNAP under standard high-yield reaction conditions were greater than that of T7 RNAP (**Table S4**) with combined error rates of 1.4 ± 0.3 x 10^-4^ and 1.3 ± 0.3 x 10^-4^ errors/base, respectively (**Fig. 2A, Table S5**). For ψ-modified RNA1/RNA5 and RNA2 the error rate was observed to be 3.1 ± 0.1 x 10^-4^ and 3.3 ± 0.7 x 10^-4^ errors/base, respectively. The total combined error rates of m1ψ-incorporated RNA1/RNA5 and RNA2 were both 2.5 ± 0 x 10^-4^ errors/base. Similar to T7 RNAP, combined error rates followed the same trend—ψ-modified RNAs demonstrating highest error rates as compared to m1ψ-modified RNAs and uridine-containing RNAs. (**Figs. 1A and 2A, Tables S2 and S5**).

**Figure 2.**
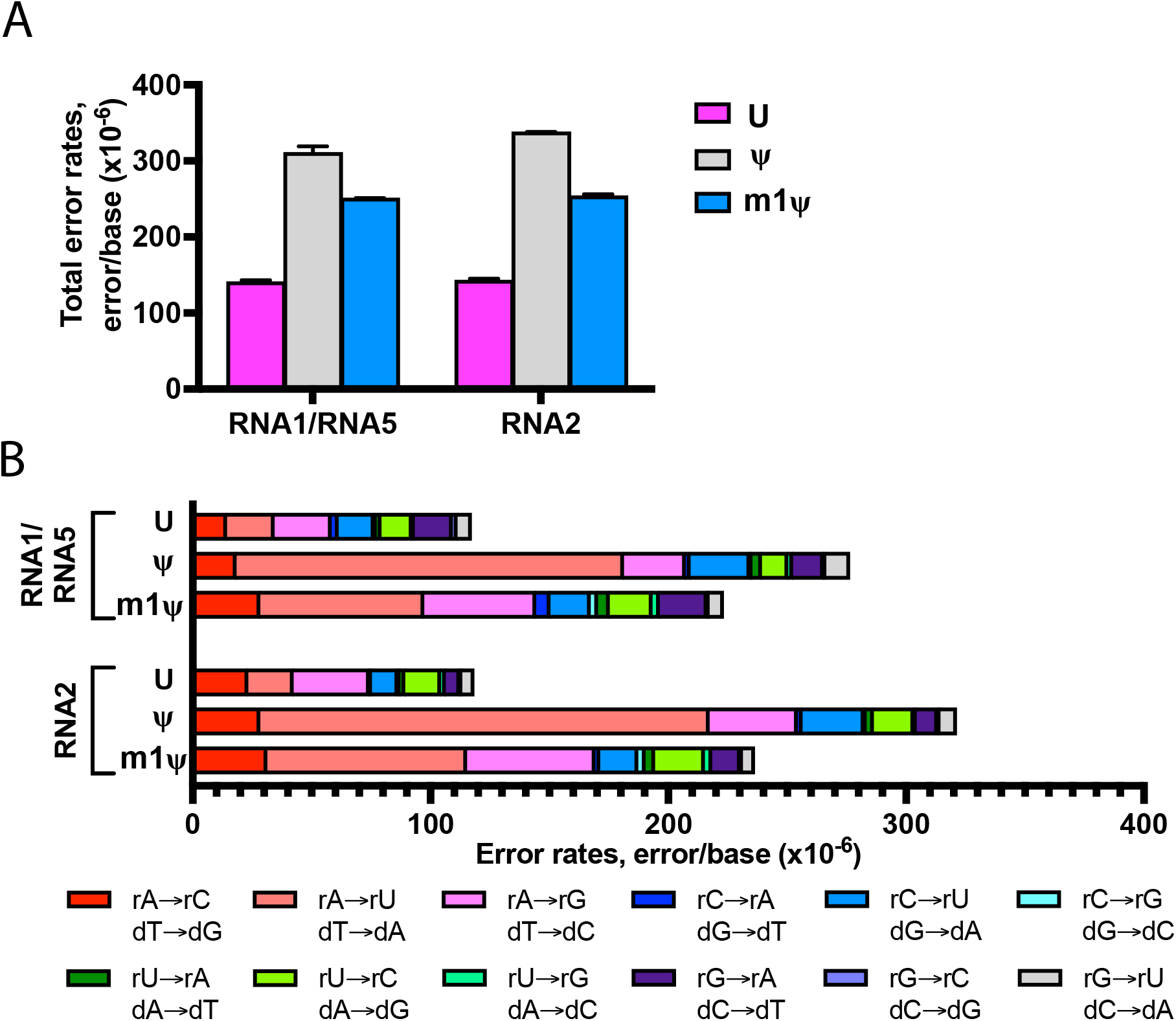
Uridine, ψ, and m1ψ are incorporated by SP6 RNA polymerase with varying error rates during *in vitro* transcription. The RNA templates synthesized with SP6 RNA polymerase were converted into cDNA with ProtoScript II reverse transcriptase followed by library preparation and sequencing. First-strand errors represented here stem from the combination of RNA polymerase and reverse transcriptase errors (**A**). RNA1 and RNA5 are artificial RNA sequences that have been permutated to include every four-base combination and RNA2 is a functional mRNA encoding Cypridina luciferase. (**B**) Base substitution error profile observed for unmodified and modified RNA sequences are represented. Colors indicate specific substitution errors observed.

Furthermore, base substitution errors were observed to be the most predominant error type ranging between 84% to 96% of total errors with SP6 RNAP (**Table S5**). In unmodified RNAs synthesized with SP6 RNAPs, the rA→rG/dT→dC substitution was observed to be the predominant error (**Fig. 2B and Table S6**). However, the substitution profiles of m1ψ- or ψ-modified RNA demonstrated a preponderance of rA→rU/dT→dA substitutions as observed with m1ψ- or ψ-modified RNAs synthesized with T7 RNAP (**Fig. 2B and Table S6**). Compared to the error rates of unmodified RNA, ψ-modified RNAs had seven-to nine-fold increase in rA→rU/dT→dA substitution in RNA1/RNA5 and RNA2, respectively. For the m1ψ-modified RNAs, the combined error rates for RNA1/RNA5 and RNA2 sequences were two-to three-fold higher than unmodified RNA. Furthermore, the sequence context analysis of the rA→rU substitutions observed in RNAs synthesized with SP6 RNAP demonstrated enrichment of a 5’ rC for both unmodified and modified RNAs (**Fig. S6**). Additionally, the substitution sites were distributed throughout the length of the transcript as observed in RNAs synthesized with T7 RNAP (**Fig. S7**).

### Fidelity of uridine incorporation is not dependent on the yield of the *in vitro* transcription reaction

For synthetic mRNA-based applications, high yield of RNA from the *in vitro* transcription reaction is desirable and reactions are typically performed with high concentrations of rNTPs. The recommended high-yield rNTP concentrations are different for T7 RNAP (40 mM rNTP) and SP6 RNAP (20 mM rNTP). In order to ensure that the differences in combined error observed for T7 RNAP and SP6 RNAP are not due to differences in the rNTP concentrations in the reactions, we performed *in vitro* transcription reactions with T7 under low rNTP reaction conditions with either 20 mM or 10 mM rNTP. The total combined error as well as the base substitution errors observed in unmodified RNA1/RNA5 when reactions were performed with T7 RNAP under low rNTP (20 or 10 mM) reaction conditions were comparable to that observed with high rNTP (40 mM) reaction conditions (**Figs. 1A and 2A, Tables S2, S4, S5, and S7**) suggesting that the overall rNTP concentration in the reaction does not affect fidelity of uridine incorporation and the differences in error rates observed with T7 and SP6 RNAP are not due to differences in the reaction conditions. Furthermore, the total combined error as well as the base substitution error profile observed in m1ψ- and ψ-modified RNAs were also unaltered under low rNTP reaction conditions (data not shown).

### Altering the rNTP composition during *in vitro* transcription reduces combined error rate and the predominant rA-to-rU substitution error

Our data demonstrates that m1ψ- and ψ-modified RNAs have increased combined errors compared to unmodified RNAs and the increased rA→rU substitution errors account for most of the misincorporations that are observed (**Figs. 1B and 2B**). Furthermore, the increased rA→rU substitution errors observed in RNAs synthesized with SP6 RNAPs suggest that the predominant rA→rU substitution might occur during *in vitro* transcription where the uridine analogs are misincorporated with higher frequency. In order to test this hypothesis as well as to reduce the rA→rU substitution during *in vitro* transcription reactions with uridine analogs, we decided to manipulate the rUTP concentration in the reaction with the idea that balancing the rUTP in the *in vitro* transcription reaction to match the nucleotide composition of the RNA sequence might result in reduced rA→rU substitution errors and consequently the total errors observed during *in vitro* transcription.

Combining sequence optimization of the synthetic RNA with incorporation of uridine modifications, specifically m1ψ and ψ, in synthetic mRNA-based vaccines and therapeutics is becoming a common practice. In addition, depleting the uridine content in the synthetic mRNA by sequence optimization has been demonstrated to reduce the immunogenicity of the synthetic molecules [28, 32]. In order to test the effect of reduced rUTP in the reactions, we generated a template for uridine-depleted randomized sequence (RNA7) that yields a final RNA base composition of 30.9% A, 33.4% C, 30.1% G, and 5.5% U. First, as a control, we analyzed the error rates when *in vitro* transcription reactions were performed under standard high-yield rNTP condition where all the rNTPs are added equally to a final concentration of 40 mM (represented as equal in **Fig. 3** and **Table S8**). The uridine-depleted RNA7 transcribed with T7 RNAP with equal molar unmodified rNTPs had a total combined error rate 6.4 ± 0.7 x 10^-5^ errors/base, while the error rates of ψ- and m1ψ-incorporating transcripts were observed to be 1.9 ± 0.2 x 10^-4^ and 9.5 ± 2.7 x 10^-5^ errors/base, respectively (**Fig. 3A and Table S8**). For the rUTP optimized reactions, rNTPs proportional to the template sequence, i.e., 12.4 mM (31%) rATP, 13.2 mM (33%) rCTP, 12 mM (30%) rGTP and 2.4 mM (6%) rUTP, were used. No difference in the RNA yield from the *in vitro* transcription reactions were observed when the rNTP levels were altered (**Table S9**). The total combined error rate of unmodified RNA was observed to be 5.2 ± 0.4 x 10^-5^ errors/base and that of m1ψ-incorporating RNA was 6.6 ± 0.5 x 10^-5^ errors/base (**Table S8**). Noticeably, the total error rate of ψ-modified RNA was 7.6 ± 0.6 x 10^-5^ errors/base, about two-fold reduced as compared to equal molar rNTP condition (1.9 ± 0.2 x 10^-4^ errors/base).

**Figure 3.**
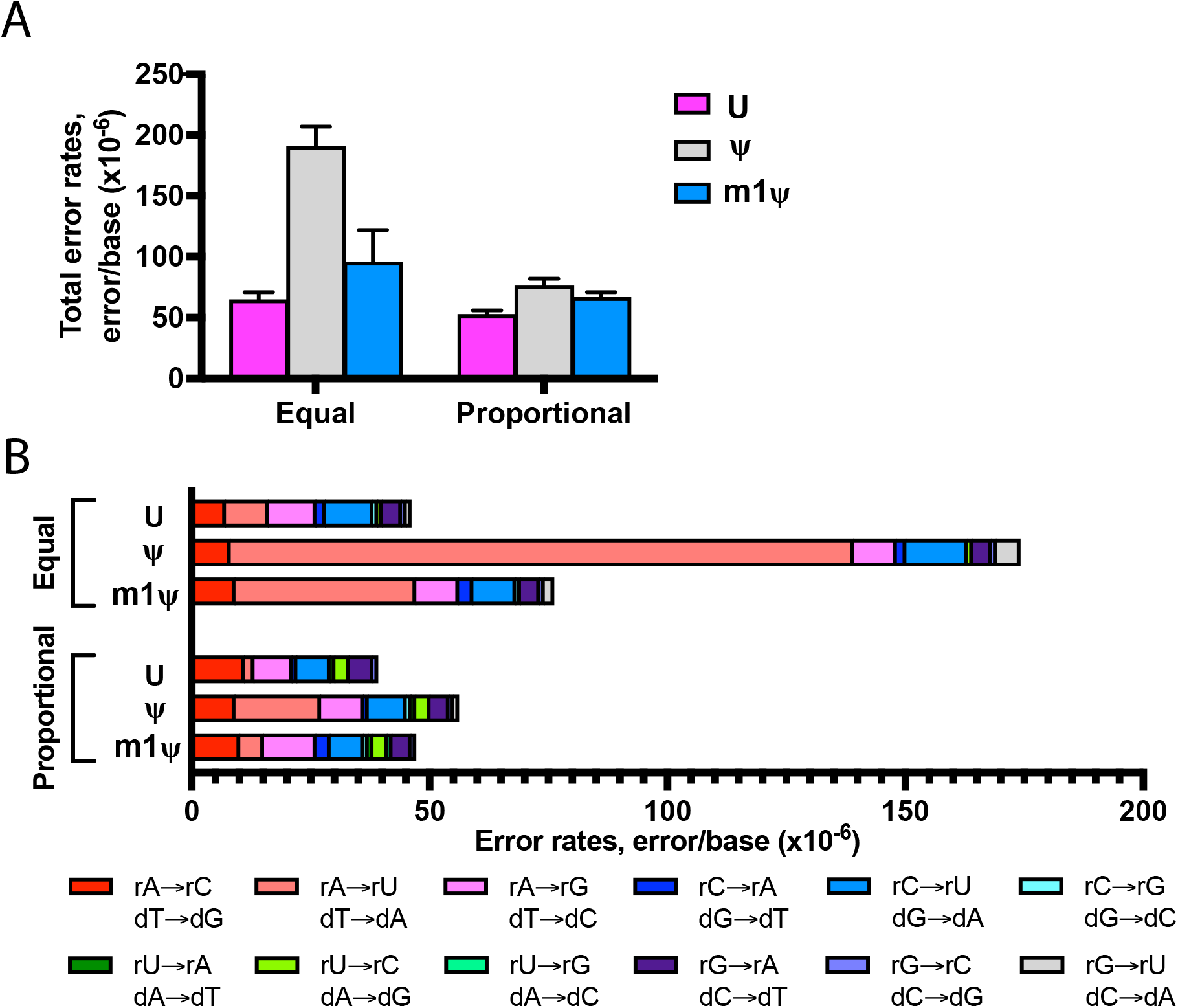
*In vitro* transcription error profile of T7 RNA polymerase can be modulated by altering ribonucleotide composition in the reaction. Uridine depleted (6% uridine content) synthetic RNA was transcribed with T7 RNA polymerase and reactions were performed with either equal molar rNTPs or rNTPs proportional to the template sequence composition. (**A**) First-strand errors represented here stem from the combination of RNA polymerase and reverse transcriptase errors. (**B**) Base substitution error profile observed for unmodified and modified RNA sequences from reactions the two different rNTP reaction conditions are represented. Colors indicate specific substitution errors observed.

Under standard equal molar rNTP reaction conditions, the substitution error profile of the uridine-depleted RNA7 sequence resembled RNA1/RNA5, RNA2 and RNA6, with rA→rU/dT→dA substitution demonstrating the most significant change when modified uridine was used in the reaction (**Fig. 3B and Table S10**). For m1ψ-modified RNA, the rA→rU/dT→dA substitution was increased three-fold as compared to the unmodified RNA and this was even more pronounced in the ψ-modified RNA with an increase of 14-fold over unmodified RNA. Interestingly, when the rNTP concentrations were altered to match the nucleotide content of the sequence, the rA→rU substitution error rates were lowered significantly as compared to that observed under standard reaction condition (equal rNTP conditions). For unmodified RNA, a four-fold reduction in rA→rU substitution was observed when rUTP concentration was altered to match the template sequence (**Fig. 3B and Table S10**). Similarly, six- and seven-fold reductions were observed for ψ-modified and m1ψ-modified RNA, respectively. Furthermore, no significant change in any of the other substitution errors were observed when the rNTP concentrations were altered (**Fig. 3B and Table S10**). The comparison of the two different RNA synthesis workflows with RNA7 demonstrates that lowering the rUTP concentration to match the nucleotide sequence of the RNA indeed reduced the rA→rU substitution error during *in vitro* transcription reaction and further provides support that the rA→rU substitutions observed with the uridine modifications are errors observed during *in vitro* transcription. Furthermore, to ensure that the effect of altering the rNTP concentration to reduce the rA→rU substitution is observed with different sequences, we generated two other templates for uridine-depleted randomized sequences, RNA8 (29.7% A, 32.0% C, 31.7% G, and 6.6% T) and RNA9 (30.2% A, 30.4% C, 27.1% G, and 12.3% T). Similar to what we observed with RNA7, reduced rA→rU substitution errors were observed for modified and unmodified RNAs (**Fig. S8; Tables S11 and S12**) when the rNTP concentrations were altered to match the sequence of the RNA.

In light of our observation that the fidelity of T7 RNAP can be modulated by adjusting the rNTP ratio in the reaction, we investigated if the same holds true for SP6 RNAP where the rA→rU substitution is also the most predominant substitution error in modified RNAs. We used T-depleted randomized sequence template (RNA7) that consists of 31% A, 33% C, 30% G, and 6% T and reactions were performed with either equimolar rNTPs (5 mM each) or rNTPs matched to the nucleotide sequence [6.2 mM (31%) rATP, 6.6 mM (33%) rCTP, 6 mM (30%) rGTP and 1.2 mM (6%) rUTP]. In presence of proportional rNTP concentrations, the total combined error rate in unmodified RNA7 was reduced to 1.3 ± 0 x 10^-4^ errors/base, one-fold decrease from 2.8 ± 0.9 x 10^-4^ errors/base in presence of equal molar unmodified rNTP (**Fig. S9A; Table S13**). Similarly, for m1ψ-modified reaction, the total combined error rate was reduced two-fold (from 4.1 ± 1.0 x 10^-4^ errors/base to 1.5 ± 0 x 10^-4^ errors/base). As observed with T7 RNAP, ψ-modified RNA had the most pronounced (three-fold) reduction in total combined error rate (from 6.0 ± 2.1 x 10^-4^ errors/base to 1.7 ± 0 x 10^-4^ errors/base) when the rNTP ratios were altered to match the sequence of RNA7. Furthermore, the reduction in total combined error observed in proportional rNTP reaction conditions is attributable specifically to reduction in the rA→rU substitution errors (**Fig. S9B; Table S14**).

U-depletion of the RNA sequence is not a viable alternative for all sequences. Furthermore, the extent of U-depletion is dependent on the sequence since it requires change in the U-content without altering the codon. We investigated a corollary approach where we hypothesized that increasing the rATP concentrations in the reaction might also reduce the rA→rU substitutions in the *in vitro* transcription reactions. We performed *in vitro* transcription of RNA1/RNA5 with excess rATPs (20 mM rATP with 10 mM of other rNTPs or 16 mM rATP with 8mM of other rNTPs). When 20mM rATP was used in the reactions, the total combined error rates of unmodified, ψ- and m1ψ-modified RNA1/RNA5 were observed to be 4.3 ± 0.1 x 10^-5^ errors/base, 5.1 ± 0.1 x 10^-5^ errors/base and 3.4 ± 0.1 x 10^-5^ errors/base, respectively (**Fig. S10A and Table S15**). A two-fold decrease in total combined error rates was observed for m1ψ-modified and ψ-modified RNAs and a 1.5-fold for unmodified RNAs (**Fig. 1 and Table S2, S16**) when the rATP concentration was increased with respect to the other rNTPs. As hypothesized, the substitution error profile of reactions performed with excess rATP showed rA→rU/dT→dA substitution in the modified RNA changed the most compared to unmodified RNA (**Fig. S10B and Table S16**). Similarly, when 16mM rATP was used in the *in vitro* transcription reactions, the total combined error rates of ψ-modified RNA1/RNA5 was observed to be 6.4 ± 0.1 x 10^-5^ errors/base in contrast to 11.0 ± 0.2 x 10^-4^ with standard rNTP conditions (**Fig. S10A and Table S17**).

Taken together, these results demonstrate that the rA→rU substitution that is observed when *in vitro* transcription reactions are performed with uridine analogs, stems from substitution errors during *in vitro* transcription with T7 and SP6 RNAP. Furthermore, the fidelity of the uridine analog incorporation can be increased by either lowering the rUTP concentration to match the nucleotide composition of the synthetic RNA sequence or increasing the rATP concentration in the reaction without compromising the yield from the reaction.

## Discussion

The use of synthetic mRNA-based vaccines and therapeutics have been evaluated for over three decades. However, the successful implementation of this modality has been hindered due to the instability of the synthetic mRNA molecule, its recognition by the cellular immune receptors, as well as the lack of an efficient delivery vehicle [1, 2]. Even though the inherent immunostimulatory profile of the synthetic mRNA together with the expression of an antigen from the synthetic mRNA has allowed for effective immune activation as well as antigen presentation [4, 5], the use of this modality for therapeutics requires the synthetic molecule to be devoid of any such unwanted response. As synthetic mRNAs get evaluated as therapeutics, it is important that the synthetic mRNAs are generated with high fidelity and are devoid of by-products that may result in unwanted immune responses. The fidelity with which modifications are incorporated in the mRNAs can affect the efficacy of the drug substance. It is becoming increasingly apparent that the enzymatic synthesis process used to generate mRNA molecules contribute significantly to the immune response observed *in vivo* [33, 34]. Rationalized design of the synthetic mRNAs, *in vitro* transcription reaction engineering, together with downstream processing of the synthetic mRNA preparations have helped ameliorate some of these effects [12, 35]. Combining chemical modifications of the synthetic mRNA with downstream purification of the mRNA preparation has become the standard for achieving efficient expression from the synthetic molecules while overcoming the immune responses [9, 36, 37]. Among the chemical modifications that are routinely introduced in synthetic mRNAs, ψ and m1ψ are favored for their ability to suppress an immune response as well as the ability to increase the translation from the mRNAs [6–8, 10–14, 18].

The current approaches to introduce modifications in synthetic mRNAs involve replacing a canonical nucleotide with a modified analog during *in vitro* transcription by ssRNAPs. T7 and SP6 RNAPs are the most commonly used RNA polymerases for *in vitro* transcription. Both T7 and SP6 have their own promoter specificities and it is also well known that they result in heterogeneous RNA populations [31, 38, 39]. However, whether or not these two RNA polymerases incorporate modified nucleotides with similar fidelities has not been tested till date. Our data demonstrates that the combined error rates in RNAs synthesized with SP6 RNAP were up to two-fold higher than those observed with T7 RNAP (**Figs. 1 and 2; Tables S2, S4 and S5**). On the other hand, T3 RNAP which has 82% sequence identity to T7 exhibited error rates comparable to that of T7 (**Table S4**). That said, it is interesting to note that the error profiles of uridine-modified RNA for both T7 and SP6 polymerases were similar. For both RNAPs, the incorporation of ψ or m1ψ was more error prone than canonical uridine incorporation (**Figs. 1A and 2A; Tables S2 and S5**). However, when uridine analogs were present in the reaction, both T7 and SP6 RNAPs had a very similar substitution error profile with rA→rU substitution being the predominant error observed during *in vitro* transcription (**Figs 1 and 2; Tables S3 and S6**). Available crystal structure capturing T7 RNAP during transcription elongation, as well as biochemical assays and molecular dynamics simulation, have identified several key residues that contribute to transcription fidelity [40–42]. For example, Tyrosine 639 is known to be critical for the pre-insertion of correct nucleotides opposite to the template DNA strand [42]. Since the differences in the total combined errors between T7 and SP6 RNAPs were observed for the same template sequences, and since base pairing between incoming rNTP and DNA template is the same, the fidelity differences of the two enzymes must be associated with differences in the protein sequence, e.g. residues which function to ensure correct base pairing. Interestingly, Tyr639 is conserved in both T7 and SP6 RNAPs as well as among other bacteriophage ssRNAPs, but neighboring amino acids are not. Future studies to elucidate the role of residues outside of the active site on fidelity of modified nucleotide incorporation will be instrumental in designing high-fidelity *in vitro* transcription systems. In addition to targeted amino acid replacements in T7 or SP6 RNAP, determining the error rates of homologous ssRNAPs will reveal the sequence determinants of transcriptional fidelity within this protein family. Furthermore, it will also be of interest to identify novel ssRNAPs that have a distinct substitution error profile than that of T7 RNAP and might be a better RNAP candidate for introduction of uridine analogs.

Even though the combined error rates observed for the same set of RNA sequences were significantly different between T7 and SP6 RNAPs, there were few similarities. First, the fidelity with which the analogs were incorporated, followed the same trend — uridine>m1ψ>ψ — with m1ψ-modified RNA exhibiting lower total errors and rA→rU/dT→dA substitution than ψ-modified RNA (**Figs 1 and 2; Tables S3 and S6**). Second, the difference in fidelity between uridine-containing RNAs and ψ/m1ψ-modified RNAs is mainly attributable to a higher rA→rU/dT→dA substitution error (**Figs. 1 and 2; Tables S3 and S6**). Based on our data from multiple templates and reaction conditions, we demonstrate that both ψ and m1ψ have a higher propensity to mis-pair with dT in the template DNA during *in vitro* transcription. Biophysical studies to measure the melting temperatures (Tm) of synthetic RNA duplexes containing either uridine, ψ or m1ψ, have shown that both ψ- or m1ψ- containing duplexes have a higher T_m_ than uridine-containing duplexes [13]. Increased base pairing and stacking has been postulated to contribute to the increased T_m_ observed [28, 43–45]. One of the critical features of ψ and m1ψ is the C5-C’1 bond that enables rotation between the sugar moiety and the nucleobase, which, in contrast to canonical uridine, could provide improved base pairing and stacking. As compared to uridine and m1ψ, ψ contains an extra hydrogen bond donor group (N1H) that imparts a universal base character to ψ. Hence, ψ can not only pair A but can also wobble basepair with G, U, or C [42]. On the other hand, m1ψ has a methyl group in the N1-position and therefore does not have the extra hydrogen bond donor and therefore wobble pairing with other nucleotides is not favored. It is plausible that the lower error rates observed in m1ψ-containing RNAs is due to the lack of the extra hydrogen bond donor [45]. Additionally, it is likely that the methyl group in the N1 position of m1ψ may alter the polarity of the C2 position and make it less favorable to pair with dT in the DNA template. It is also worth noting that stability difference between RNA duplex containing four consecutive ψ’s and m1ψ’s was not distinguishable [13]. Stability measurements in which a single rψTP or rm1ψTP pairs with DNA template to mimic the errors observed during *in vitro* transcription will likely be more relevant to distinguish the base-pairing differences we observe in this study. Not much is understood about the pairing of rψ or rm1ψ with DNA, but there is precedence for RNA-RNA interactions in presence of ψ and m1ψ which might help us understand the increased error rates observed in presence of the modified analogs [13, 44–46].

Another aspect of rationalized mRNA design that has been gaining traction has been to reduce the uridine composition without altering the amino acid sequence of the protein encoded from the synthetic mRNA [28, 32]. Uridine depletion in Cas9 mRNA sequence demonstrated a reduction of innate immune response and an increase in Cas9 activity [32]. Comirnaty and Spikevax sequences consist of 19% and 15% uridine, respectively, as compared to the wild-type spike protein sequence that has 33% uridine in the sequence [27, 47]. Often, rationalized design of the synthetic mRNA molecule is further combined with reaction optimization such as altering the rNTP concentrations in the reaction to optimize the RNA yields from the reactions as well as reduction of dsRNA byproducts [12, 18]. Our data demonstrates that combining uridine depletion of the RNA sequence with altering the rNTP composition of the reaction, reduces the rA→rU substitutions that are introduced during *in vitro* transcription (**Figs. 3B, S8, S9, and S10; Tables S10, S12, S14-S16**). Future studies aimed at comparing the outcome from these modified processes (uridine depletion with altered rNTP composition to generate high-fidelity synthetic mRNA with reduced dsRNA byproducts) will provide further insights into ways to optimize the efficacy from the synthetic mRNA drug substance.

Our observation that the prevalent rA→rU substitutions introduced during *in vitro* transcription can be reduced by balancing the rNTPs in the reaction suggests a few possible mechanisms for introduction of these substitution errors. By limiting the rψTP amounts or competing with excess rATP, the rA→rU substitution can be altered to reduce the error rates without affecting any of the other substitution errors (**Figs. 3, S8, S9, S10; Tables S8, S10-S16**). One of the possible reasons for this could be, that these mispairing events occur in presence of excess rNTPs in the reaction. This is further supported by the fact that in experiments where the template sequence was intentionally biased to have reduced uridine (RNA7), adding equal molar rNTPs in the reaction (10mM each) led to an overall increased error profile than that observed in templates that had comparable nucleotide usage (RNA2). Interestingly, the nucleotide composition of RNA1 and RNA5 have equal representation of all the four nucleotides and we still observed prevalent rA→rU substitutions when equal molar rNTPs were added in the reaction (**Figs 1B and 2B, Tables S3 and S6**). These reactions were performed with 10 mM rNTP each. It is possible that not all the rNTPs in the reaction are utilized by T7 RNAP there is always excess of rNTPs in these high-yield rection conditions. Future studies to dissect out if the misincorporations are more prevalent in the early *vs* late stage of the *in vitro* transcription process will provide further insight into the events leading to uridine analog misincorporation. It can also be envisioned that *in vitro* transcription systems that limit the initial rNTP concentration but allow for a steady, optimized rNTP feeding mechanism might further help improve the fidelity of nucleotide incorporation and be instrumental in generating high-fidelity synthetic mRNAs.

In order to achieve high-fidelity RNA products, it is desirable to understand the rules of nucleotide incorporation so that if there are sequences that are more error prone, they can be omitted from the synthetic mRNA during design. However, in this study, the comparison of multiple sequence contexts demonstrated that other than T7 RNAP-incorporated ψ-modified RNAs that demonstrated slight sequence context preference, there is no strong correlation between the sequence context of the DNA template and the substitution errors observed under any condition tested (**Figs. S4 and S6**). Furthermore, the errors observed were distributed throughout of the length of RNA did not affect the misincorporation events that were observed when *in vitro* transcription was performed with either T7 RNAP or SP6 RNAP and when uridine analogs were used (**Figs. S5 and S7**).

Our results demonstrate that the presence of ψ and m1ψ in the *in vitro* transcription reactions result in higher base substitution errors in the modified RNAs. A critical aspect to consider is how these errors might affect the efficacy of the synthetic mRNA drug substance and how much of these error-prone molecules can be tolerated *in vivo* without any adverse effect. Because *in vitro* transcribed mRNAs are modified throughout the body of the mRNA, it is also critical to consider if these mRNAs are faithfully translated in the cell. We have previously demonstrated that in human embryonic kidney cell, low frequency translation elongation miscoding events are observed from ψ-containing mRNAs due to altered tRNA selection in ψ-containing codons [22]. For m1ψ-substituted RNAs, it has been shown that translation initiation and ribosome transit is altered *in vivo* [14]. A complete understanding of what errors are incorporated during the RNA synthesis process and how that further affects the identity of the protein synthesized is a timely question given that therapeutic applications would require repeat dosing of the mRNAs and would also require expression of the protein of choice. To be able to predict the best outcome from these synthetic molecules, it is critical that we understand where variability comes from and to be able to define the rules to avoid these variabilities.

## Acknowledgements

The authors would like to thank Dillon Nye, Guy Birkenmeier, George Tzertzinis for comments on the manuscript and Rick Morgan, Sean Maguire, Alexey Fomenkov for help with the PacBio sequencing.

## Author contribution

B.R and T.H.C conceived and designed the experiments; T.H.C and N.D carried out the experiments; V.P and T.H.C analyzed the data; J.L.O and B.R contributed reagents; B.R. and T.H.C wrote the manuscript with input from all authors.

