## Supplementary information for "Improving the fidelity of uridine analog incorporation during *in vitro* transcription"

#### Supplementary figure legends

**Supplemental Figure 1. T7 and SP6 RNA polymerases incorporate U, ψ, and m1ψ into full-length RNAs**. (**A**) Representative bioanalyzer traces of RNA1 and RNA2 transcribed with uridine analogs showing synthesis of the full-length RNA and the integrity of the RNA. (**B**) Representative UHPLC traces for RNA1. The purified RNA digested with a nucleoside digestion mix was analyzed. (**C**) Incorporation of uridine analogs in RNA1 and RNA2 assessed using UHPLC/MS. All the ψ- and m1ψ-modified transcripts had higher than 99.6% incorporation of modified uridine. Data reported are an average of three independent experiments.

**Supplemental Figure 2. T7 RNA polymerase synthesizes run-off products and run-off products with non-templated additions in presence of U, ψ, and m1ψ**. Short 30-nt RNA (RNA3) and 60-nt RNA (RNA4) were transcribed *in vitro* and assessed using intact mass spectrometry. From the top four abundant masses, the RNAs that were either transcribed as expected or with one or two non-templated nucleotide additions are highlighted in the orange box as run-off transcripts.

**Supplemental Figure 3. Schematic illustration of workflow to measure combined fidelity of RNA polymerases and ProtoScript II reverse transcriptase**. (**A**) Linear dsDNA served as a template to synthesize RNA with unmodified and modified uridine by T7 or SP6 RNA polymerases. The transcript was reverse transcribed into first strand and second strand cDNA by ProtoScript II reverse transcriptase. The dsDNA was ligated with SMRTbell adaptor and subjected for sequencing on the PacBio system. Substitution errors observed (for example a rA→rU/dT→dA) can occur either during *in vitro* transcription or reverse transcription. (**B**) During *in vitro* transcription, RNA polymerase might mis-incorporate an rU (red) in place of rA. The correct incorporation of rA (red) and the resulting cDNA were shown. (**C**) During the first strand cDNA synthesis, reverse transcriptase might mis-incorporate a dT (red) to the position of dA. The correct incorporation of dA and the resulting cDNA were shown.

**Supplemental Figure 4. rA→rU/dT→dA substitution errors observed in ψ–containing RNAs have sequence context preference when reactions are performed with T7 RNA polymerase**. The sequence surrounding the rA→rU/dT→dA substitution sites were used to generate the sequence logo. The data was generated from four different RNAs: synthetic RNA1 and RNA5 with sequence permutation to include every four-base combination, and two functional mRNAs, RNA2 and RNA6. Seven nucleotides upstream and downstream of the substitution site were examined and for simplicity the immediate sequence upstream and downstream of the substitution site is represented. The numbers of sites plotted were summarized in Table S18.

**Supplemental Figure 5. rA→rU/dT→dA substitution errors are observed along the length of the RNA with no preference for a specific position in the transcript when reactions are performed with T7 RNA polymerase**. Four RNAs were assessed: synthetic RNA1 and RNA5 with sequence permutation to include every four-base combination, and two functional mRNAs, RNA2 and RNA6.

**Supplemental Figure 6. rA→rU/dT→dA substitution errors observed in RNAs synthesized with SP6 RNA polymerase do not have sequence context preference**. The sequence surrounding the rA→rU/dT→dA substitution sites were used to generate the sequence logo. The data was generated from four different RNAs: synthetic RNA1 and RNA5 with sequence permutation to include every four-base combination, and a functional mRNA, RNA2. Seven nucleotides upstream and downstream of the substitution site were examined and for simplicity the immediate sequence upstream and downstream of the substitution site is represented. The numbers of sites plotted were summarized in Table S18.

**Supplemental Figure 7. rA→rU/dT→dA substitution errors are observed along the length of the RNA with no preference for a specific position in the transcript when reactions are performed with SP6 RNA polymerase**. Three RNAs were assessed: synthetic RNA1 and RNA5 with sequence permutation to include every 4-base combination, and a functional mRNA, RNA2.

**Supplemental Figure 8. *In vitro* transcription error profile of T7 RNA polymerase can be modulated by altering ribonucleotide composition in the reaction**. Uridine-depleted synthetic RNA sequence RNA8 (8% uridine content) and RNA9 (12% uridine content) were transcribed with T7 RNA polymerase and reactions were performed with either equal molar rNTPs or rNTPs proportional to the template sequence composition. (**A**) First-strand errors represented here stem from the combination of RNA polymerase and reverse transcriptase errors. (**B**) Base substitution error profile observed for unmodified and modified RNA sequences from reactions the two different rNTP reaction conditions are represented. Colors indicate specific substitution errors observed.

**Supplemental Figure 9. *In vitro* transcription error profile of SP6 RNA polymerase can be modulated by altering ribonucleotide composition in the reaction**. Uridine-depleted synthetic RNA sequence RNA7 (6% uridine content) was transcribed with SP6 RNA polymerase and reactions were performed with either equal molar rNTPs or rNTPs proportional to the template sequence composition. (**A**) First-strand errors resulting from the combined errors from SP6 RNA polymerase and Protoscript II reverse transcriptase are represented. (**B**) Base substitution error profile observed for unmodified and modified RNA sequences from reactions the two different rNTP reaction conditions are represented. Colors indicate specific substitution errors observed.

**Supplemental Figure 10. *In vitro* transcription error profile of T7 RNA polymerase can be modulated by altering rATP composition in the reaction.**

*In vitro* transcription reactions were performed with T7 RNA polymerase using 20 mM rATP (instead of 10 mM rATP in standard IVT conditions). RNA1 and RNA5 are artificial RNA sequence that has been permutated to include every four-base combination. (**A**) Total combined errors of RNA polymerase and reverse transcriptase for transcripts consisting of uridine analogs. (**B**) Base substitution error profile observed for unmodified and modified RNA sequences from reactions with excess rATP are represented. Colors indicate specific substitution errors observed.

#### Supplementary Tables

| Platform | Total error rate (×$10^{-6}$, errors/base) | Substitution (%) | Deletion (%) | Insertion (%) | Total sequenced bases | Reference |
|---|---|---|---|---|---|---|
| RSII | 56 ± 8 | 71 | 19 | 10 | 30,868,961 | [1] |
| RSII | 41 ± 4 | 80 | 11 | 9 | 12,024,831 | This study |
| Sequel | 64 ± 4 | 73 | 12 | 15 | 7,820,043 | This study |

**Supplemental Table 1**. Comparison of combined error rates for unmodified RNA1/RNA5 synthesized with T7 RNA polymerase measured using either RSII or Sequel platform.

| **Template** | **Base** | **Total error rate ($\times 10^{-6}$, errors/base)** | **Substitution (%)** | **Deletion (%)** | **Insertion (%)** | **Total sequenced bases** |
|---|---|---|---|---|---|---|
| RNA1/RNA5 | U | 64 ± 4 | 73 | 12 | 15 | 7,820,043 |
| RNA1/RNA5 | ψ | 110 ± 20 | 91 | 4 | 5 | 2,764,788 |
| RNA1/RNA5 | m1ψ | 80 ± 3 | 84 | 9 | 7 | 3,097,951 |
| RNA2 | U | 61 ± 0 | 89 | 7 | 4 | 29,699,388 |
| RNA2 | ψ | 144 ± 32 | 96 | 3 | 2 | 10,396,843 |
| RNA2 | m1ψ | 83 ± 4 | 89 | 8 | 3 | 906,212 |
| RNA6 | U | 47 ± 1 | 91 | 6 | 3 | 7,522,053 |
| RNA6 | ψ | 125 ± 2 | 96 | 4 | 0 | 1,552,716 |
| RNA6 | m1ψ | 69 ± 4 | 91 | 7 | 2 | 5,733,742 |

**Supplemental Table 2**. Total and specific error rates of first strand cDNA observed in RNA1/RNA5, RNA2 and RNA6 synthesized with T7 RNA polymerase.

| RNA | Uridine analog | Substitution error rate ($\times 10^{-6}$, errors/base) | | | | | | | | | | | |
|---|---|---|---|---|---|---|---|---|---|---|---|---|---|
| | | rA→rC dT→dG | rA→rU dT→dA | rA→rG dT→dC | rC→rA dG→dT | rC→rU dG→dA | rC→rG dG→dC | rU→rA dA→dT | rU→rC dA→dG | rU→rG dA→dC | rG→rA dC→dT | rG→rC dC→dG | rG→rU dC→dA |
| RNA1/RNA5 | U | 4 | 5 | 7 | 2 | 7 | 1 | 1 | 8 | 2 | 8 | 1 | 2 |
| | ψ | 4 | 58 | 4 | 8 | 7 | 0 | 3 | 4 | 1 | 5 | 0 | 4 |
| | m1ψ | 8 | 15 | 10 | 5 | 7 | 0 | 2 | 7 | 1 | 9 | 1 | 1 |
| | **Relative fold change (ψ to U)** | **0** | **11** | **0** | **3** | **0** | **-1** | **2** | **-1** | **-1** | **0** | **-1** | **1** |
| | **Relative fold change (m1ψ to U)** | **1** | **2** | **0** | **2** | **0** | **-1** | **1** | **0** | **-1** | **0** | **0** | **-1** |
| RNA2 | U | 11 | 10 | 11 | 1 | 8 | 0 | 1 | 7 | 1 | 2 | 1 | 1 |
| | ψ | 7 | 98 | 9 | 2 | 7 | 1 | 1 | 4 | 1 | 3 | 1 | 4 |
| | m1ψ | 8 | 23 | 14 | 6 | 9 | 0 | 4 | 4 | 2 | 3 | 0 | 0 |
| | **Relative fold change (ψ to U)** | **0** | **9** | **0** | **1** | **0** | **N/A** | **0** | **0** | **0** | **1** | **0** | **3** |
| | **Relative fold change (m1ψ to U)** | **0** | **1** | **0** | **5** | **0** | **N/A** | **3** | **0** | **1** | **1** | **-1** | **-1** |
| RNA6 | U | 4 | 6 | 6 | 1 | 11 | 0 | 1 | 5 | 1 | 5 | 1 | 1 |
| | ψ | 3 | 79 | 7 | 1 | 14 | 1 | 0 | 3 | 1 | 6 | 2 | 3 |
| | m1ψ | 7 | 22 | 5 | 3 | 9 | 1 | 2 | 5 | 1 | 5 | 1 | 1 |
| | **Relative fold change (ψ to U)** | **0** | **12** | **0** | **0** | **0** | **N/A** | **-1** | **0** | **0** | **0** | **1** | **2** |
| | **Relative fold change (m1ψ to U)** | **1** | **3** | **0** | **2** | **0** | **N/A** | **1** | **0** | **0** | **0** | **0** | **0** |

**Supplemental Table 3.** Substitution error rates of first strand cDNA observed in RNA1/RNA5, RNA2 and RNA6 synthesized with T7 RNA polymerase. The relative fold change was calculated for each substitution as (M—U) / U, where M is the substitution rate on modified RNA and U is the substitution rate on unmodified RNA. N/A, not applicable, as the error rates were rounded to zero in unmodified RNA.

| **RNA polymerase** | **Total error rate ($\times10^{-6}$, errors/base)** | **Substitution (%)** | **Deletion (%)** | **Insertion (%)** | **Total sequenced bases** |
|---|---|---|---|---|---|
| T3 | 68 ± 10 | 71 | 9 | 20 | 10,981,293 |
| T7 | 64 ± 4 | 73 | 12 | 15 | 7,820,043 |
| SP6 | 140 ± 3 | 84 | 6 | 10 | 5,463,829 |

**Supplemental Table 4**. Total and specific error rates of first strand cDNA observed in unmodified RNA1/RNA5 synthesized with T3, T7 and SP6 RNA polymerases.

| Template | Base | Total error rate ($\times10^{-6}$, errors/base) | Substitution (%) | Deletion (%) | Insertion (%) | Total sequenced bases |
|---|---|---|---|---|---|---|
| RNA1/RNA5 | U | 140 ± 3 | 84 | 6 | 10 | 5,463,829 |
| RNA1/RNA5 | ψ | 310 ± 9 | 90 | 3 | 7 | 4,047,319 |
| RNA1/RNA5 | m1ψ | 250 ± 1 | 89 | 4 | 7 | 4,116,031 |
| RNA2 | U | 132 ± 3 | 90 | 7 | 3 | 33,183,812 |
| RNA2 | ψ | 337 ± 1 | 96 | 3 | 1 | 7,005,019 |
| RNA2 | m1ψ | 253 ± 3 | 94 | 5 | 2 | 34,593,081 |

**Supplemental Table 5**. Total and specific error rates of first strand cDNA observed in RNA1/RNA5, RNA2 and RNA6 synthesized with SP6 RNA polymerase.

| RNA | Uridine analog | Substitution error rate (×$10^{-6}$, errors/base) | | | | | | | | | | | |
|---|---|---|---|---|---|---|---|---|---|---|---|---|---|
| | | **rA→rC dT→dG** | **rA→rU dT→dA** | **rA→rG dT→dC** | **rC→rA dG→dT** | **rC→rU dG→dA** | **rC→rG dG→dC** | **rU→rA dA→dT** | **rU→rC dA→dG** | **rU→rG dA→dC** | **rG→rA dC→dT** | **rG→rC dC→dG** | **rG→rU dC→dA** |
| RNA1/RNA5 | U | 14 | 20 | 24 | 3 | 15 | 1 | 2 | 13 | 1 | 16 | 2 | 6 |
| | ψ | 18 | 163 | 26 | 2 | 25 | 1 | 4 | 11 | 2 | 13 | 1 | 10 |
| | m1ψ | 28 | 69 | 47 | 6 | 17 | 3 | 5 | 18 | 3 | 20 | 1 | 6 |
| | **Relative fold change (ψ to U)** | **0** | **7** | **0** | **0** | **1** | **0** | **1** | **0** | **1** | **0** | **-1** | **1** |
| | **Relative fold change (m1ψ to U)** | **1** | **2** | **1** | **1** | **0** | **2** | **2** | **0** | **2** | **0** | **-1** | **0** |
| RNA2 | U | 23 | 19 | 32 | 1 | 11 | 1 | 2 | 15 | 2 | 6 | 1 | 5 |
| | ψ | 28 | 189 | 37 | 2 | 26 | 1 | 3 | 17 | 1 | 9 | 1 | 7 |
| | m1ψ | 31 | 84 | 54 | 2 | 16 | 3 | 4 | 21 | 3 | 12 | 1 | 5 |
| | **Relative fold change (ψ to U)** | **0** | **9** | **0** | **1** | **1** | **0** | **1** | **0** | **-1** | **1** | **0** | **0** |
| | **Relative fold change (m1ψ to U)** | **0** | **3** | **1** | **1** | **0** | **2** | **1** | **0** | **1** | **1** | **0** | **0** |

**Supplemental Table 6.** Substitution error rates of first strand cDNA observed in RNA1/RNA5 and RNA2 synthesized with SP6 RNA polymerase. The relative fold change was calculated for each substitution as (M—U) / U, where M is the substitution rate on modified RNA and U is the substitution rate on unmodified RNA.

| **rNTP (total)** | **Base** | **Total error rate ($\times 10^{-6}$, errors/base)** | **Substitution (%)** | **Deletion (%)** | **Insertion (%)** | **Total sequenced bases** |
|---|---|---|---|---|---|---|
| 40 mM | U | 64 ± 4 | 73 | 12 | 15 | 7,820,043 |
| 20 mM | U | 64 ± 2 | 77 | 10 | 13 | 16,735,875 |
| 10 mM | U | 66 ± 9 | 78 | 10 | 12 | 9,406,619 |

**Supplemental table S7.** Total and specific error rates of first strand cDNA observed in RNA1/RNA5 synthesized with T7 RNA polymerase under varying rNTP concentrations.

| **rNTP** | **Base** | **Total error rate ($\times10^{-6}$, errors/base)** | **Substitution (%)** | **Deletion (%)** | **Insertion (%)** | **Total sequenced bases** |
|---|---|---|---|---|---|---|
| Equal | U | 64 ± 7 | 71 | 16 | 13 | 7,554,213 |
| Equal | ψ | 190 ± 17 | 91 | 4 | 5 | 16,176,735 |
| Equal | m1ψ | 95 ± 27 | 82 | 9 | 9 | 11,640,523 |
| Proportional | U | 52 ± 4 | 76 | 13 | 12 | 15,834,035 |
| Proportional | ψ | 76 ± 6 | 74 | 12 | 14 | 18,366,763 |
| Proportional | m1ψ | 66 ± 5 | 73 | 13 | 14 | 21,837,801 |

**Supplemental Table 8.** Total and specific error rates of first strand cDNA observed in uridine-depleted RNA7 synthesized with T7 RNA polymerase.

| rNTP | Base | Yield (μg) |
|---|---|---|
| Equal | U | 80 |
| Equal | ψ | 68 |
| Equal | m1ψ | 50 |
| Proportional | U | 44 |
| Proportional | ψ | 70 |
| Proportional | m1ψ | 64 |

**Supplemental Table 9.** Representative yield from *in vitro* transcription reactions performed with T7 RNA polymerase with either equal molar rNTPs or rNTPs proportional to the template sequence (RNA7).

| rNTP | Uridine analog | Substitution error rate (×10$^{-6}$, errors/base) | | | | | | | | | | | |
|---|---|---|---|---|---|---|---|---|---|---|---|---|---|
| | | **rA→rC dT→dG** | **rA→rU dT→dA** | **rA→rG dT→dC** | **rC→rA dG→dT** | **rC→rU dG→dA** | **rC→rG dG→dC** | **rU→rA dA→dT** | **rU→rC dA→d G** | **rU→rG dA→dC** | **rG→rA dC→dT** | **rG→rC dC→dG** | **rG→rU dC→dA** |
| Equal | U | 7 | 9 | 10 | 2 | 10 | 1 | 0 | 1 | 0 | 4 | 1 | 1 |
| | ψ | 8 | 131 | 9 | 2 | 13 | 0 | 0 | 1 | 0 | 4 | 1 | 5 |
| | m1ψ | 9 | 38 | 9 | 3 | 9 | 1 | 0 | 0 | 0 | 4 | 1 | 2 |
| | **Relative fold change (ψ to U)** | **0** | **14** | **0** | **0** | **0** | **-1** | **N/A** | **0** | **N/A** | **0** | **0** | **4** |
| | **Relative fold change (m1ψ to U)** | **0** | **3** | **0** | **1** | **0** | **0** | **N/A** | **-1** | **N/A** | **0** | **0** | **1** |
| Proportional | U | 11 | 2 | 8 | 1 | 7 | 0 | 1 | 3 | 0 | 5 | 1 | 0 |
| | ψ | 9 | 18 | 9 | 1 | 8 | 1 | 1 | 3 | 0 | 4 | 1 | 1 |
| | m1ψ | 10 | 5 | 11 | 3 | 7 | 1 | 1 | 3 | 1 | 4 | 1 | 0 |
| | **Relative fold change (ψ to U)** | **0** | **8** | **0** | **0** | **0** | **N/A** | **0** | **0** | **N/A** | **0** | **0** | **N/A** |
| | **Relative fold change (m1ψ to U)** | **0** | **2** | **0** | **2** | **0** | **N/A** | **0** | **0** | **N/A** | **0** | **0** | **N/A** |

**Supplemental Table 10.** Substitution error rates of first strand cDNA observed in uridine-depleted RNA7 synthesized with T7 RNA polymerase. The relative fold change was calculated for each substitution as (M—U) / U, where M is the substitution rate on modified RNA and U is the substitution rate on unmodified RNA. N/A, not applicable, as the error rates were rounded to zero in unmodified RNA.

| RNA | rNTP | Base | Total error rate ($\times 10^{-6}$, errors/base) | Substitution (%) | Deletion (%) | Insertion (%) | Total sequenced bases |
|---|---|---|---|---|---|---|---|
| RNA8 | Equal | U | 75 ± 9 | 63 | 34 | 3 | 11,196,978 |
| RNA8 | Equal | ψ | 220 ± 48 | 86 | 12 | 2 | 10,182,271 |
| RNA8 | Equal | m1ψ | 150 ± 17 | 84 | 12 | 3 | 21,302,209 |
| RNA8 | Proportional | U | 91 ± 15 | 53 | 43 | 4 | 15,890,495 |
| RNA8 | Proportional | ψ | 84 ± 3 | 70 | 25 | 5 | 8,418,903 |
| RNA8 | Proportional | m1ψ | 75 ± 6 | 74 | 24 | 2 | 3,174,159 |
| RNA9 | Equal | U | 62 ± 11 | 86 | 7 | 7 | 10,349,048 |
| RNA9 | Equal | ψ | 140 ± 12 | 96 | 2 | 2 | 7,784,454 |
| RNA9 | Equal | m1ψ | 80 ± 11 | 90 | 7 | 3 | 10,476,693 |
| RNA9 | Proportional | U | 50 ± 4 | 83 | 13 | 5 | 7,135,004 |
| RNA9 | Proportional | ψ | 75 ± 9 | 93 | 4 | 3 | 14,246,294 |
| RNA9 | Proportional | m1ψ | 60 ± 2 | 83 | 11 | 6 | 17,833,482 |

**Supplemental Table 11.** Total and specific error rates of first strand cDNA observed in uridine-depleted RNA8 and RNA9 synthesized with T7 RNA polymerase.

| RNA | rNTP | Uridine analog | Substitution error rate (×10$^{-6}$, errors/base) | | | | | | | | | | | |
|---|---|---|---|---|---|---|---|---|---|---|---|---|---|---|
| | | | rA→rC dT→dG | rA→rU dT→dA | rA→rG dT→dC | rC→rA dG→dT | rC→rU dG→dA | rC→rG dG→dC | rU→rA dA→dT | rU→rC dA→dG | rU→rG dA→dC | rG→rA dC→dT | rG→rC dC→dG | rG→rU dC→dA |
| RNA8 | Equal | U | 7 | 11 | 7 | 2 | 10 | 1 | 0 | 1 | 0 | 4 | 1 | 2 |
| | | ψ | 6 | 134 | 9 | 4 | 15 | 1 | 0 | 1 | 0 | 7 | 2 | 7 |
| | | m1ψ | 8 | 49 | 10 | 15 | 17 | 2 | 1 | 1 | 1 | 12 | 2 | 5 |
| | | **Relative fold change (ψ to U)** | **0** | **11** | **0** | **1** | **1** | **0** | **N/A** | **0** | **N/A** | **1** | **1** | **3** |
| | | **Relative fold change (m1ψ to U)** | **0** | **3** | **0** | **7** | **1** | **1** | **N/A** | **0** | **N/A** | **2** | **1** | **2** |
| | Proportional | U | 10 | 2 | 11 | 3 | 8 | 2 | 1 | 2 | 1 | 7 | 2 | 1 |
| | | ψ | 8 | 20 | 10 | 2 | 8 | 2 | 0 | 2 | 1 | 5 | 1 | 1 |
| | | m1ψ | 8 | 8 | 9 | 4 | 8 | 1 | 1 | 4 | 0 | 7 | 2 | 1 |
| | | **Relative fold change (ψ to U)** | **0** | **9** | **0** | **0** | **0** | **0** | **-1** | **0** | **0** | **0** | **-1** | **0** |
| | | **Relative fold change (m1ψ to U)** | **0** | **3** | **0** | **0** | **0** | **-1** | **0** | **1** | **-1** | **0** | **0** | **0** |
| RNA9 | Equal | U | 11 | 8 | 11 | 2 | 8 | 1 | 0 | 3 | 1 | 6 | 1 | 1 |
| | | ψ | 10 | 95 | 8 | 2 | 11 | 1 | 0 | 1 | 1 | 4 | 1 | 2 |
| | | m1ψ | 9 | 27 | 12 | 4 | 9 | 1 | 1 | 1 | 1 | 5 | 0 | 2 |
| | | **Relative fold change (ψ to U)** | **0** | **11** | **0** | **0** | **0** | **0** | **N/A** | **-1** | **0** | **0** | **0** | **1** |
| | | **Relative fold change (m1ψ to U)** | **0** | **2** | **0** | **1** | **0** | **0** | **N/A** | **-1** | **0** | **0** | **-1** | **1** |
| | Proportional | U | 9 | 2 | 9 | 1 | 6 | 1 | 1 | 4 | 1 | 5 | 1 | 1 |
| | | ψ | 8 | 35 | 6 | 2 | 7 | 1 | 1 | 2 | 1 | 4 | 1 | 2 |
| | | m1ψ | 8 | 9 | 10 | 4 | 6 | 2 | 1 | 3 | 1 | 4 | 1 | 1 |
| | | **Relative fold change (ψ to U)** | **0** | **17** | **0** | **1** | **0** | **0** | **0** | **-1** | **0** | **0** | **0** | **1** |
| | | **Relative fold change (m1ψ to U)** | **0** | **4** | **0** | **3** | **0** | **1** | **0** | **0** | **0** | **0** | **0** | **0** |

**Supplemental Table 12.** Total and specific error rates of first strand cDNA observed in uridine-depleted RNA8 and RNA9 synthesized with T7 RNA polymerase. The relative fold change was calculated for each substitution as (M—U) / U, where M is the substitution rate on modified RNA and U is the substitution rate on unmodified RNA. N/A, not applicable, as the error rates were rounded to zero in unmodified RNA.

| rNTP | Base | Total error rate ($\times 10^{-6}$, errors/base) | Substitution (%) | Deletion (%) | Insertion (%) | Total sequenced bases |
|---|---|---|---|---|---|---|
| Equal | U | 281 ± 86 | 92 | 4 | 4 | 10,587,337 |
| Equal | ψ | 597 ± 210 | 96 | 2 | 2 | 17,735,541 |
| Equal | m1ψ | 407 ± 95 | 94 | 3 | 3 | 7,454,925 |
| Proportional | U | 131 ± 1 | 87 | 9 | 5 | 12,731,671 |
| Proportional | ψ | 172 ± 1 | 91 | 5 | 4 | 26,186,535 |
| Proportional | m1ψ | 154 ± 2 | 88 | 6 | 6 | 7,985,264 |

**Supplemental Table 13.** Total and specific error rates of first strand cDNA observed in uridine-depleted RNA7 synthesized with SP6 RNA polymerase.

| rNTP | Uridine analog | Substitution error rate ($\times10^{-6}$, errors/base) | | | | | | | | | | | |
|---|---|---|---|---|---|---|---|---|---|---|---|---|---|
| | | **rA→rC dT→dG** | **rA→rU dT→dA** | **rA→rG dT→dC** | **rC→rA dG→dT** | **rC→rU dG→dA** | **rC→rG dG→dC** | **rU→rA dA→dT** | **rU→rC dA→dG** | **rU→rG dA→dC** | **rG→rA dC→dT** | **rG→rC dC→dG** | **rG→rU dC→dA** |
| Equal | U | 39 | 77 | 47 | 4 | 50 | 2 | 1 | 3 | 0 | 17 | 2 | 18 |
| | ψ | 29 | 406 | 39 | 3 | 55 | 3 | 1 | 2 | 0 | 15 | 2 | 21 |
| | m1ψ | 46 | 200 | 59 | 5 | 30 | 3 | 2 | 3 | 1 | 19 | 2 | 13 |
| | **Relative fold change (ψ to U)** | **0** | **4** | **0** | **0** | **0** | **1** | **0** | **0** | **N/A** | **0** | **0** | **0** |
| | **Relative fold change (m1ψ to U)** | **0** | **2** | **0** | **0** | **0** | **1** | **1** | **0** | **N/A** | **0** | **0** | **0** |
| Proportional | U | 32 | 5 | 27 | 3 | 7 | 1 | 3 | 17 | 2 | 13 | 2 | 2 |
| | ψ | 28 | 42 | 33 | 3 | 11 | 2 | 3 | 14 | 2 | 15 | 2 | 3 |
| | m1ψ | 29 | 14 | 37 | 4 | 9 | 3 | 3 | 16 | 2 | 17 | 2 | 1 |
| | **Relative fold change (ψ to U)** | **0** | **7** | **0** | **0** | **1** | **1** | **0** | **0** | **0** | **0** | **0** | **1** |
| | **Relative fold change (m1ψ to U)** | **0** | **2** | **0** | **0** | **0** | **2** | **0** | **0** | **0** | **0** | **0** | **-1** |

**Supplemental Table 14.** Substitution error rates of first strand cDNA observed in uridine-depleted RNA7 synthesized by SP6 RNA polymerase. The fold change between modified and unmodified RNA is highlighted in red. The relative fold change was calculated for each substitution as (M—U) / U, where M is the substitution rate on modified RNA and U is the substitution rate on unmodified RNA. N/A, not applicable, as the error rates were rounded to zero in unmodified RNA.

| rATP | Base | Total error rate ($\times 10^{-6}$, errors/base) | Substitution (%) | Deletion (%) | Insertion (%) | Total sequenced bases |
|---|---|---|---|---|---|---|
| 10 mM | U | 64 ± 4 | 73 | 12 | 15 | 7,820,043 |
| 10 mM | ψ | 110 ± 20 | 91 | 4 | 5 | 2,764,788 |
| 10 mM | m1ψ | 80 ± 3 | 84 | 9 | 7 | 3,097,951 |
| 20 mM | U | 43 ± 1 | 71 | 18 | 2 | 19,570,384 |
| 20 mM | ψ | 51 ± 1 | 82 | 9 | 9 | 21,978,065 |
| 20 mM | m1ψ | 34 ± 1 | 81 | 14 | 5 | 13,742,187 |

**Supplemental Table 15.** Total error rates of first strand cDNA observed in RNA1/RNA5 synthesized with T7 RNA polymerase in presence of 10 or 20 mM rATP

| rATP | Uridine analog | Substitution error rate (×$10^{-6}$, errors/base) | | | | | | | | | | | |
|---|---|---|---|---|---|---|---|---|---|---|---|---|---|
| | | **rA→rC dT→dG** | **rA→rU dT→dA** | **rA→rG dT→dC** | **rC→rA dG→dT** | **rC→rU dG→dA** | **rC→rG dG→dC** | **rU→rA dA→dT** | **rU→rC dA→dG** | **rU→rG dA→dC** | **rG→rA dC→dT** | **rG→rC dC→dG** | **rG→r[illegible] dC→d[illegible]** |
| 10 mM | U | 4 | 5 | 7 | 2 | 7 | 1 | 1 | 8 | 2 | 8 | 1 | 2 |
| | ψ | 4 | 58 | 4 | 8 | 7 | 0 | 3 | 4 | 1 | 5 | 0 | 4 |
| | **Relative fold change (ψ to U)** | **0** | **11** | **0** | **3** | **0** | **-1** | **2** | **-1** | **-1** | **0** | **-1** | **1** |
| 16 mM | U | 2 | 1 | 4 | 1 | 6 | 0 | 4 | 8 | 1 | 5 | 1 | 2 |
| | ψ | 3 | 20 | 6 | 2 | 9 | 0 | 2 | 5 | 1 | 6 | 1 | 4 |
| | **Relative fold change (ψ to U)** | **1** | **19** | **1** | **1** | **1** | **N/A** | **-1** | **0** | **0** | **0** | **0** | **1** |

**Supplemental Table 16.** Substitution error rates of first strand cDNA observed in RNA1/RNA5 synthesized with T7 RNA polymerase in presence of 10 mM or 16 mM rATP. The relative fold change was calculated for each substitution as (M—U) / U, where M is the substitution rate on modified RNA and U is the substitution rate on unmodified RNA. N/A, not applicable, as the error rates were rounded to zero in unmodified RNA. not applicable, as the error rates were rounded to zero in unmodified RNA.

| rATP | Base | Total error rate (×$10^{-6}$, errors/base) | Substitution (%) | Deletion (%) | Insertion (%) | Total sequenced bases |
|---|---|---|---|---|---|---|
| 10 mM | U | 64 ± 4 | 73 | 12 | 15 | 7,820,043 |
| 10 mM | ψ | 110 ± 20 | 91 | 4 | 5 | 2,764,788 |
| 16 mM | U | 45 ± 5 | 78 | 10 | 12 | 9,146,473 |
| 16 mM | ψ | 64 ± 1 | 92 | 4 | 4 | 11,588,053 |

**Supplemental Table 17.** Total error rates of first strand cDNA observed in RNA1/RNA5 synthesized with T7 RNA polymerase in presence of 10 or 16 mM rATP.

#### Total reads

| me | RNA | Uridine analog | Substitution | Deletion | nsertion | rA→rC dT→dG | rA→rU dT→dA | rA→rG dT→dC | rC→rA dG→dT | rC→rU dG→dA | rC→rG dG→dC | rU→rA dA→dT | rU→rC dA→dG | rU→rG dA→dC | rG→rA dC→dT | rG→rC dC→dG | rG→rU dC→dA |
|---|---|---|---|---|---|---|---|---|---|---|---|---|---|---|---|---|---|
| T7 RNAP | RNA1/ RNA5 | U | 368 | 62 | 73 | 30 | 36 | 53 | 19 | 55 | 5 | 7 | 59 | 18 | 63 | 8 | 15 |
| | | $\psi$ | 265 | 11 | 15 | 10 | 159 | 11 | 21 | 18 | 0 | 8 | 10 | 3 | 14 | 1 | 10 |
| | | m1$\psi$ | 208 | 22 | 18 | 26 | 48 | 32 | 14 | 21 | 1 | 7 | 23 | 3 | 27 | 2 | 4 |
| | RNA2 | U | 1619 | 121 | 71 | 339 | 289 | 334 | 23 | 231 | 6 | 22 | 210 | 41 | 63 | 24 | 37 |
| | | $\psi$ | 1429 | 42 | 25 | 73 | 1018 | 94 | 18 | 75 | 10 | 8 | 39 | 14 | 30 | 13 | 37 |
| | | m1$\psi$ | 67 | 6 | 2 | 7 | 21 | 13 | 5 | 8 | 0 | 4 | 4 | 2 | 3 | 0 | 0 |
| | RNA6 | U | 322 | 20 | 12 | 33 | 45 | 44 | 9 | 84 | 3 | 9 | 37 | 5 | 41 | 5 | 7 |
| | | $\psi$ | 185 | 8 | 0 | 5 | 122 | 11 | 2 | 22 | 1 | 0 | 4 | 1 | 10 | 3 | 4 |
| | | m1$\psi$ | 362 | 28 | 7 | 42 | 125 | 31 | 19 | 52 | 4 | 11 | 26 | 6 | 31 | 7 | 8 |
| SP6 RNAP | RNA1/ RNA5 | U | 645 | 49 | 74 | 78 | 109 | 130 | 14 | 84 | 7 | 13 | 72 | 8 | 88 | 11 | 31 |
| | | $\psi$ | 1121 | 35 | 83 | 72 | 660 | 104 | 10 | 102 | 5 | 17 | 44 | 8 | 53 | 4 | 42 |
| | | m1$\psi$ | 926 | 44 | 72 | 116 | 286 | 195 | 26 | 72 | 13 | 20 | 76 | 12 | 83 | 4 | 23 |
| | RNA2 | U | 3925 | 318 | 140 | 764 | 630 | 1063 | 46 | 374 | 33 | 64 | 483 | 55 | 202 | 34 | 177 |
| | | $\psi$ | 2255 | 73 | 30 | 198 | 1322 | 262 | 17 | 185 | 5 | 23 | 116 | 10 | 62 | 7 | 48 |
| | | m1$\psi$ | 8202 | 399 | 139 | 1078 | 2920 | 1853 | 67 | 562 | 100 | 141 | 719 | 114 | 414 | 49 | 185 |

**Supplemental Table 18.** Substitutions, deletions and insertions for first strand cDNA synthesis observed in RNA1, RNA2, RNA5 and RNA6 synthesized with T7 and SP6 RNA polymerases.

| Oligonucleotide | Sequence |
|---|---|
| Forward primer for RNA1, RNA5, RNA8, RNA9 | AGAGTACACGAGTCAGGCTACAGCATC |
| Reverse primer for RNA1, RNA5, RNA8, RNA9 | TACAGTTCACGAGGACCGTCAAGA |
| Forward primer for RNA2 | CGCCACCATGAAGACCTTAA |
| Reverse primer for RNA2 | GTCTCCATGCTTTATGTAGC |
| Forward primer for RNA6 | TGAACCTGACCACCAGAACA |
| Reverse primer for RNA6 | AACTAGCAGAGGTGGTGAGT |
| Forward primer for RNA7 | AAGCCACCTAGAACAACCAC |
| Reverse primer for RNA7 | TTCGTTGGCTTGGACACTTCTG |

**Supplementary Table 19**. Oligonucleotide sequence for reverse transcription.

| Construct | Sequence |
|---|---|
| T7p_RNA1 | TAATACGACTCACTATAGGGTCTAGAAATAATTTTGTTTAACTTTAGAGTACACGAGTCA<br>GGCTACAGCATCCTCTGGTTCAGACTACTTGATTCATGTGTACCCTATATGCGAGGATAT<br>GTGTATCGTAGAAATTGTCAGGCAGTAACGTTCCGCGAGTTTTAATGGGCGCGCCATGA<br>CTCTAAGAGTGATATACCTCCTCGGTCTCGGGCCCGGGGTGTAATTAGCCCAGTTAGAC<br>ACGATCGCCCGACGTATATTGTTGCTTGGGTATCGTCGCATGCGAAGTATTGCCCAAGG<br>AGACACAACAAGCAACTTATGTTGACTCCCTTCGACCATTAAAATTTGTTAGAACGGACA<br>GAAAGGATGCGCCTTATAAATGTCCTGTGCAGTGATGAAGCGACCTCAAAACGCTTCAT<br>GATCTAACCGACTCACCTTGCCGTTCCCTCCGCGCCTTAAAACCGGCCGGTCTTGCGAA<br>AAGCGGGAAACGAGTTTACCCACGGATAGCAGGGAATGTTGCGGCTGGCTAGGGAGCA<br>TGAAGGTAGATACTCCACGGCTTACCTTTCCGGGGCTCAACATCTAGCCACAGACCTTT<br>TCGTTAAGCCCACCCCCACTGGATACTGAATCATCAGGGAACCGGACCCAACCAGTTTG<br>GGCTCGTCCAAGCTTCGGTCTCGTCCCTAAGTGCAAAGATATGGAAAGAGCAGCATAG<br>GTATATGGATTATTCTTTTACCACTCGTTTCTTACCGTAACTTACGCAATGGATCACGTG<br>CCGAGGCGGCGGTACAGCTGTTCGAAGGGCTCTGTCGGGAACGCTAACATCCAGCCGG<br>TAAATTCCAAACTAGGGAAAGGACACGCACTGAATTGAATATAGTCGTGAAGGGTGGTG<br>TAAGTCGTGCACAGCCCGCATTAAGTACTAAACAGCGTCCAATCTTGATCTACTTACGG<br>CCTGATGTTCTTCAGCACCTCCTAGCACTGGAGTACTTCGCTATCAATGAGATTAGCACT<br>TTGTACATGTCATCCAGCCCGAGTCTGGGGTCCGACAATGCGGTCGCCGATTGGTATCT<br>GCATGTAGTATTAAACGGAGCTGCCGCGGCTGCGGATTATAGTTCATGTCTTGACGGTC<br>CTCGTGAACTGTG**GTTAAC** |
| T7p_RNA2 | TAATACGACTCACTATAGGGAGACCCAAGCTTGGTACCGAGCTCGGATCCGCCACCATG<br>AAGACCTTAATTCTTGCCGTTGCATTAGTCTACTGCGCCACTGTTCATTGCCAGGACTGT<br>CCTTACGAACCTGATCCACCAAACACAGTTCCAACTTCCTGTGAAGCTAAAGAAGGAGAA<br>TGTATTGATAGCAGCTGTGGCACCTGCACGAGAGACATACTATCAGATGGACTGTGTGA<br>AAATAAACCAGGAAAAACATGTTGCCGAATGTGTCAGTATGTAATTGAATGCAGAGTAGA<br>GGCCGCAGGATGGTTTAGAACATTCTATGGAAAGAGATTCCAGTTCCAGGAACCTGGTA<br>CATACGTGTTGGGTCAAGGAACCAAGGGCGGCGACTGGAAGGTGTCCATCACCCTGGA<br>GAACCTGGATGGAACCAAGGGGGCTGTGCTGACCAAGACAAGACTGGAAGTGGCTGGA<br>GACATCATTGACATCGCTCAAGCTACTGAGAATCCCATCACTGTAAACGGTGGAGCTGAC<br>CCTATCATCGCCAACCCGTACACCATCGGCGAGGTCACCATCGCTGTTGTTGAGATGCC<br>AGGCTTCAACATCACCGTCATTGAGTTCTTCAAACTGATCGTGATCGACATCCTCGGAGG<br>AAGATCTGTAAGAATCGCCCCAGACACAGCAAACAAAGGAATGATCTCTGGCCTCTGTG<br>GAGATCTTAAAATGATGGAAGATACAGACTTCACTTCAGATCCAGAACAACTCGCTATTC<br>AGCCTAAGATCAACCAGGAGTTTGACGGTTGTCCACTCTATGGAAATCCTGATGACGTTG<br>CATACTGCAAAGGTCTTCTGGAGCCGTACAAGGACAGCTGCCGCAACCCCATCAACTTCT<br>ACTACTACACCATCTCCTGCGCCTTCGCCCGCTGTATGGGTGGAGACGAGCGAGCCTCA<br>CACGTGCTGCTTGACTACAGGGAGACGTGCGCTGCTCCCGAAACTAGAGGAACCTGCGT<br>TTTGTCTGGACATACTTTCTACGATACATTTGACAAAGCAAGATACCAATTCCAGGGTCC |

| | |
|---|---|
| | CTGCAAGGAGATTCTTATGGCCGCCGACTGTTTCTGGAACACTTGGGATGTGAAGGTTTCACACAGGAATGTTGACTCTTACACTGAAGTAGAGAAAGTACGAATCAGGAAACAATCGACTGTAGTAGAACTCATTGTTGATGGAAAACAGATTCTGGTTGGAGGAGAAGCCGTGTCCGTCCCGTACAGCTCTCAGAACACTTCCATCTACTGGCAAGATGGTGACATACTGACTACAGCCATCCTACCTGAAGCTCTGGTGGTCAAGTTCAACTTCAAGCAACTGCTCGTCGTACATATTAGAGATCCATTCGATGGTAAGACTTGCGGTATTTGCGGTAACTACAACCAGGATTTCAGTGATGATTCTTTTGATGCTGAAGGAGCCTGTGATCTGACCCCCAACCCACCGGGATGCACCGAAGAACAGAAACCTGAAGCTGAACGACTCTGCAATAGTCTCTTCGCCGGTCAAAGTGATCTTGATCAGAAATGTAACGTGTGCCACAAGCCTGACCGTGTCGAACGATGCATGTACGAGTATTGCCTGAGGGGACAACAGGGTTTCTGTGACCACGCATGGGAGTTCAAGAAAGAATGCTACATAAAGCATGGAGACACCCTAGAAGTACCAGATGAATGCAAATAG**GCGGCCGC** |
| T7p_RNA3 | CAGTAATACGACTCACTATAGGTGACATACTGACTACAGCCATCCTACCT |
| T7p_RNA4 | CAGTAATACGACTCACTATAGGAAATCCTGATGACGTTGCATACTGCAAAGGTCTTCTGGAGCCGTACAAGGACAGCTGC |
| T7p_RNA5 | TAATACGACTCACTATAGGGTCTAGAAATAATTTTGTTTAACTTTAGAGTACACGAGTCAGGCTACAGCATCTTGACACCAGAATATTATGGATTGGACGCTTCCCACTAAATGGAAGACTGTTCGGTCATAAACACTACTAGGAATTCCTCTCCAGTCATCATGTTCGATCGTCTAGCAGCAATCTCTTCCGATCGATATTTGCGCGTGACTCAGGCGAGCCCATGACAGCTTCTCCCCGTGAGAACCACGACTAGAAGTTATCTGTTGAGCTGCTAGCTTCGTGGCCCGGCCATGGTAGTAGCGGCTCACTCGCGCTAACTTTGCCTGCTCGAGAAAACGGGCGAAACACCCAGCAACACAAGCCACTTAATTTGTTGATAGATAATAAGATCAGGTTATTAGTCGCTCTGCACTTACTTTAAGTGCCAACTATGCTGTATCGGCCAGGGTGAAAACGGGTGCCGCCACTtCaGTGTGTCGGAGTCTGCTGACGGATTAGGGCACAGACGTATGGTTATATCCTAAGGTAGTGTGTCAATGTACTGGGGACAAAGTCAGTGGGCACCGCATCAGGAGTGCAACCTCCGCTAGTACCGACTCGTCAATGCTTTGAGCGATGGCTTGCGCTCCCAAATCCTTAAGCTTTTATGCATTCGGCTCTGGCCCTCAGGCCTGACCTGGAATTTCATCGGAAACGCCTTAACCGACATTACATCGACACCAAGATCCCGACGCTTCATGCGGAGACGATAGAGACTCTAACCAAGAATAAAAGGAGTAGTCCCTAATCTACTGAAACGGGGATACCTCAAATCACGGGAATGCGTTACTGACCCGCTATGTGAGGCTCGGATCACCCTCGTTCTATTGCCTTGTAATCATGGTGGGGCGGCGGAGCGGGATTAGAGGGTGTCCCTAATGTGAGTAGATCTGTAGTAATGATACGTCTCCTCAATATGAGGCGTATTGCAGGTCACAGCACAGGGAGATTTCGGCGCACCCAGCCGAGTTGCCTCCGTCGTTGTTTAGGTATATGCATAACTGCTCACGACAAATACAGCAGAGCCTACGTTGGGTTATCGAATCCTTGTGGACAAGAAGCTTCTTCATGTCTTGACGGTCCTCGTGAACTGTG**GTTAAC** |
| T7p_RNA6 | TAATACGACTCACTATAGGAGAATAAACTAGTATTCTTCTGGTCCCCACAGACTCAGAGAGAACCCGCCACCATGTTCGTGTTCCTGGTGCTGCTGCCTCTGGTGTCCAGCCAGTGTGTGAACCTGACCACCAGAACACAGCTGCCTCCAGCCTACACCAACAGCTTTACCAGAGGCGTGTACTACCCCGACAAGGTGTTCAGATCCAGCGTGCTGCACTCTACCCAGGACCTGTTCCTGCCTTTCTTCAGCAACGTGACCTGGTTCCACGCCATCCACGTGTCCGGCACCAATGGCACCAAGAGATTCGACAACCCCGTGCTGCCCTTCAACGACGGGGTGTACTTTGCCAGCACCGAGAAGTCCAACATCATCAGAGGCTGGATCTTCGGCACCACACTGGACAGCAAGACCCA |

| | |
|---|---|
| | GAGCCTGCTGATCGTGAACAACGCCACCAACGTGGTCATCAAAGTGTGCGAGTTCCAGT<br>TCTGCAACGACCCCTTCCTGGGCGTCTACTACCACAAGAACAACAAGAGCTGGATGGAA<br>AGCGAGTTCCGGGTGTACAGCAGCGCCAACAACTGCACCTTCGAGTACGTGTCCCAGCC<br>TTTCCTGATGGACCTGGAAGGCAAGCAGGGCAACTTCAAGAACCTGCGCGAGTTCGTGT<br>TTAAGAACATCGACGGCTACTTCAAGATCTACAGCAAGCACACCCCTATCAACCTCGTGC<br>GGGATCTGCCTCAGGGCTTCTCTGCTCTGGAACCCCTGGTGGATCTGCCCATCGGCATC<br>AACATCACCCGGTTTCAGACACTGCTGGCCCTGCACAGAAGCTACCTGACACCTGGCGA<br>TAGCAGCAGCGGATGGACAGCTGGTGCCGCCGCTTACTATGTGGGCTACCTGCAGCCTA<br>GAACCTTCCTGCTGAAGTACAACGAGAACGGCACCATCACCGACGCCGTGGATTGTGCT<br>CTGGATCCTCTGAGCGAGACAAAGTGCACCCTGAAGTCCTTCACCGTGGAAAAGGGCAT<br>CTACCAGACCAGCAACTTCCGGGTGCAGCCCACCGAATCCATCGTGCGGTTCCCCAATA<br>TCACCAATCTGTGCCCCTTCGGCGAGGTGTTCAATGCCACCAGATTCGCCTCTGTGTACG<br>CCTGGAACCGGAAGCGGATCAGCAATTGCGTGGCCGACTACTCCGTGCTGTACAACTCC<br>GCCAGCTTCAGCACCTTCAAGTGCTACGGCGTGTCCCCTACCAAGCTGAACGACCTGTG<br>CTTCACAAACGTGTACGCCGACAGCTTCGTGATCCGGGGAGATGAAGTGCGGCAGATTG<br>CCCCTGGACAGACAGGCAAGATCGCCGACTACAACTACAAGCTGCCCGACGACTTCACC<br>GGCTGTGTGATTGCCTGGAACAGCAACAACCTGGACTCCAAAGTCGGCGGCAACTACAA<br>TTACCTGTACCGGCTGTTCCGGAAGTCCAATCTGAAGCCCTTCGAGCGGGACATCTCCA<br>CCGAGATCTATCAGGCCGGCAGCACCCCTTGTAACGGCGTGGAAGGCTTCAACTGCTAC<br>TTCCCACTGCAGTCCTACGGCTTTCAGCCCACAAATGGCGTGGGCTATCAGCCCTACAG<br>AGTGGTGGTGCTGAGCTTCGAACTGCTGCATGCCCCTGCCACAGTGTGCGGCCCTAAGA<br>AAAGCACCAATCTCGTGAAGAACAAATGCGTGAACTTCAACTTCAACGGCCTGACCGGC<br>ACCGGCGTGCTGACAGAGAGCAACAAGAAGTTCCTGCCATTCCAGCAGTTTGGCCGGGA<br>TATCGCCGATACCACAGACGCCGTTAGAGATCCCCAGACACTGGAAATCCTGGACATCA<br>CCCCTTGCAGCTTCGGCGGAGTGTCTGTGATCACCCCTGGCACCAACACCAGCAATCAG<br>GTGGCAGTGCTGTACCAGGACGTGAACTGTACCGAAGTGCCCGTGGCCATTCACGCCGA<br>TCAGCTGACACCTACATGGCGGGTGTACTCCACCGGCAGCAATGTGTTTCAGACCAGAG<br>CCGGCTGTCTGATCGGAGCCGAGCACGTGAACAATAGCTACGAGTGCGACATCCCCATC<br>GGCGCTGGAATCTGCGCCAGCTACCAGACACAGACAAACAGCCCTCGGAGAGCCAGAA<br>GCGTGGCCAGCCAGAGCATCATTGCCTACACAATGTCTCTGGGCGCCGAGAACAGCGTG<br>GCCTACTCCAACAACTCTATCGCTATCCCCACCAACTTCACCATCAGCGTGACCACAGAG<br>ATCCTGCCTGTGTCCATGACCAAGACCAGCGTGGACTGCACCATGTACATCTGCGGCGA<br>TTCCACCGAGTGCTCCAACCTGCTGCTGCAGTACGGCAGCTTCTGCACCCAGCTGAATA<br>GAGCCCTGACAGGGATCGCCGTGGAACAGGACAAGAACACCCAAGAGGTGTTCGCCCA<br>AGTGAAGCAGATCTACAAGACCCCTCCTATCAAGGACTTCGGCGGCTTCAATTTCAGCCA<br>GATTCTGCCCGATCCTAGCAAGCCCAGCAAGCGGAGCTTCATCGAGGACCTGCTGTTCA<br>ACAAAGTGACACTGGCCGACGCCGGCTTCATCAAGCAGTATGGCGATTGTCTGGGCGAC<br>ATTGCCGCCAGGGATCTGATTTGCGCCCAGAAGTTTAACGGACTGACAGTGCTGCCTCC<br>TCTGCTGACCGATGAGATGATCGCCCAGTACACATCTGCCCTGCTGGCCGGCACAATCA<br>CAAGCGGCTGGACATTTGGAGCAGGCGCCGCTCTGCAGATCCCCTTTGCTATGCAGATG<br>GCCTACCGGTTCAACGGCATCGGAGTGACCCAGAATGTGCTGTACGAGAACCAGAAGCT<br>GATCGCCAACCAGTTCAACAGCGCCATCGGCAAGATCCAGGACAGCCTGAGCAGCACAG |

| | |
|---|---|
| | CAAGCGCCCTGGGAAAGCTGCAGGACGTGGTCAACCAGAATGCCCAGGCACTGAACAC<br>CCTGGTCAAGCAGCTGTCCTCCAACTTCGGCGCCATCAGCTCTGTGCTGAACGATATCCT<br>GAGCAGACTGGACCCTCCTGAGGCCGAGGTGCAGATCGACAGACTGATCACAGGCAGA<br>CTGCAGAGCCTCCAGACATACGTGACCCAGCAGCTGATCAGAGCCGCCGAGATTAGAGC<br>CTCTGCCAATCTGGCCGCCACCAAGATGTCTGAGTGTGTGCTGGGCCAGAGCAAGAGAG<br>TGGACTTTTGCGGCAAGGGCTACCACCTGATGAGCTTCCCTCAGTCTGCCCCTCACGGC<br>GTGGTGTTTCTGCACGTGACATATGTGCCCGCTCAAGAGAAGAATTTCACCACCGCTCCA<br>GCCATCTGCCACGACGGCAAAGCCCACTTTCCTAGAGAAGGCGTGTTCGTGTCCAACGG<br>CACCCATTGGTTCGTGACACAGCGGAACTTCTACGAGCCCCAGATCATCACCACCGACA<br>ACACCTTCGTGTCTGGCAACTGCGACGTCGTGATCGGCATTGTGAACAATACCGTGTAC<br>GACCCTCTGCAGCCCGAGCTGGACAGCTTCAAAGAGGAACTGGACAAGTACTTTAAGAA<br>CCACACAAGCCCCGACGTGGACCTGGGCGATATCAGCGGAATCAATGCCAGCGTCGTG<br>AACATCCAGAAAGAGATCGACCGGCTGAACGAGGTGGCCAAGAATCTGAACGAGAGCC<br>TGATCGACCTGCAAGAACTGGGGAAGTACGAGCAGTACATCAAGTGGCCCTGGTACATC<br>TGGCTGGGCTTTATCGCCGGACTGATTGCCATCGTGATGGTCACAATCATGCTGTGTTGC<br>ATGACCAGCTGCTGTAGCTGCCTGAAGGGCTGTTGTAGCTGTGGCAGCTGCTGCAAGTT<br>CGACGAGGACGATTCTGAGCCCGTGCTGAAGGGCGTGAAACTGCACTACACATGATGAC<br>TCGAGCTGGTACTGCATGCACGCAATCCTAGCTGCCCCTTTCCCGTCCTGGGTACCCCG<br>AGTCTCCCCCGACCTCGGGTCCCAGGTATGCTCCCACCTCCACCTGCCCCACTCACCAC<br>CTCTGCTAGTTCCAGACACCTCCCAAGCACGCAGCAATGCAGCTCAAAACGCTTAGCCTA<br>GCCACACCCCCACGGGAAACAGCAGTGATTAACCTTTAGCAATAAACGAAAGTTTAACTA<br>AGCTATACTAACCCCAGGGTTGGTCAATTTCGTGCCAGCCACACCCTGGACCTAGCGCG<br>GCCG**GCTAGC** |
| T7p_RNA7 | TAATACGACTCACTATAGGGTCTAGAGCGTCGGCAAGCCACCTAGAACAACCACGGCAC<br>GCGAACCAACAGCGGCAACGCCGAACGGCCACGCGACGAACTGAGCACCACCACAAGA<br>CCAAGCCAACGAGGAACACCGAACAACCGGTCACTGAACGCAACGGCCTAGGCGCGCA<br>CACCAATAAGACGGATCAAGCCAGCGCATGACTGCGCCGGACAACGCCACGGACGACG<br>AACAGGTCCACCGGAGTACGACTATGCCATACCAGCAGGACAGCGGAGGCGACCAACA<br>ACAACAGGAGCGGCAGCCGGCAGACGAATTGGACCGACCGGACATGGCCACATGCGAT<br>TGAGCCGGAACCGCGCGCACTTGACAGTCAGCAACCGGCTCCAACAGTCCGGCACCAT<br>GAGAGAACCGGCAAGCCGGCCTTGCCACAGGAACGCAGAGAGCGGAGCAACACAAGCC<br>GACGCCAAGCCAACCGCGAACCAGCCAGGCGCCGCCGGCAACAGGAAGGTCGCACGA<br>GCCAACATATGGACCGGCCGGAACACTAAGGACCAAGAGGCCACTGGCGCGGAACGGA<br>GAACCAGGCACCAAGCAAGGAACAGAGAGGCATAGGAACCGAAGGCGCCATAGCCGGC<br>GAGACGACCAGGAACGAACTGACAGCCAACGCTAGAGGAACGTGACGATGGACGAACG<br>TAGACACGGCGGCGCAACGCCGCGGCGGAGTCCGGAACCGGCGCTAAGGTCCACACG<br>GCAGAACCGGCCAGGCCGAGGCAACCGCCGCCACACGACCACAGCAGCCAGAAGGATA<br>GCGCGCGGCCATAACAGGAACAGCAAGGACAGGCGCCACGCAGGACCGCAACTGAGCA<br>ACCGGCAACCACCTCCACCGAAGCCGCGAAGCAGAGGCCGACGACGGCACACGCGCCA<br>TGAGCGGCACAACCAGCCAGAAGGCCAACGCGCCAAGGCGACCGAAGCCAACTACAGA<br>AGTGTCCAAGCCAACGAAGCCAACCGGCAGAAGAGAGCAGG**GTTAAC** |

| | |
|---|---|
| T7p_RNA8 | TAATACGACTCACTATAGGGTCTAGAAATAATTTTGTTTAACTTTAGAGTACACGAGTCAG<br>GCTACAGCATCCTCCAGGCAGCCACACAACGGAGGCAAGAGCGGCAAGCGCAACAACG<br>CGCTCCGCGACGAGCGCGCAGAGTATCGGCAGATTCCACCGGACGGAGCACGAGGACG<br>AAGCGACCGGTCCACACGGCTGGAGGACGCAGCGCCAAGCTCGGTGCCGACCGAAGCC<br>GAGCGAGAGGAACGCCGAGAAGCAAGCCGCGACTCCACAGAGCCGCCAGCCGAAGAA<br>GACGCACGCCAGGAGGAACAGGAAGCGAGCGGAAGGTCAACGCACCAAGGCCAGCAG<br>GACTAAGCCGGCAGACGGCGGCCGCCACCACCAAGCGAACAATGATACACGAGAACGG<br>ACCGGCAACATGGACACGGCGTAGTCCACAGAGCCACAAGGCAGTGGTGCGAACCGGC<br>AACGGAGAACAATACCGGAAGAGCACAACAGAGAGGAGCTGGCCGGCACACGAGCGCG<br>ACGCGGCACCGGCGCGCCACACCAGCCGAAGGCAGCCAGCACCAACGAGCGCGGCGC<br>ACCGACTGACGGCGAGAACAGACCGAAGGCCGGCGGCCACCACGGACGCGGACCAGA<br>AGACCGGATGGCAGGCGACAACGACGGCTCGCCTCCAACAAGAAGCGGCGGACCGGC<br>GGAAGCGACAGGCGCCGCACGTGAACACAACAACTGCACGGCATGCGGCGAACGGCCA<br>AGCGAGACGACGCGCTCCAAGAGCCAACGATAGAATGCTACGGCAACCGAGCAGAGGC<br>AGAGCGCCAGCGCAACACACCAGAAGAACGCGACGAGAGAGACGGACGCCGGCGCGG<br>ACCATTGGAGGCGGAGAACCGACAACCGCAATGCGGAGATCAAGGCGGACCGAACGAC<br>AACAGCAGAGCCACCAACTGCGCCGTGGTCGCAACCAACCTAACCAGCAACAACAGGCA<br>CCGCCGCGGCGACACAGGACGAGGCCGCCGAACAACGATAACGCGACACCAGGCACAC<br>ACACGAACCGCGGAAGGACCACAGCGACAGCCATTCATGTCTTGACGGTCCTCGTGAAC<br>TGTG**GTTAAC** |
| T7p_RNA9 | TAATACGACTCACTATAGGGTCTAGAAATAATTTTGTTTAACTTTAGAGTACACGAGTCAG<br>GCTACAGCATCCTCACACACCAATGATCCAATAACCTCGAAGGCGACACCGCATGTCGC<br>CACCAGCCGTGAACGACGGAAGAATAACCGCAGCGAATACGGACATCAAGAGGCACAG<br>AACCTACGAAGCAACACGAACGCACACCATAACTTCAGCGGCAACGAAGCCGACTGAGG<br>TCCGGCCGGCAACCAAGGCCGAAGTTCACTAAGCCTCGTAACGGCAGCCGTATAGGACC<br>TCGCCGAGCACCAGACGACACCAAGAGAAGGCAGCGACCAACTGCGTCGGCCGGCGAA<br>CTGCCGAAGCTAAGGCACAGACCGGTCGCCAGGCCGCAGACCGCGGAACCAACAAGAG<br>GCAGCGCTAGGCCATCTCAGCACCACCAAGACATTGATATTAACAGAACGCGGAATCAA<br>GCGTACGATACCAACGACAAGTGTGACGAACGTGCGTCTGAATGGCACATAAGGTCGCC<br>GGTAGCGAACGCCGAACCGAACCACCTACCGCAACCAACCGACACCATAGGATGTCACA<br>GACGACTCGCAACGGCTAGACCTTACCAACACAGGAACATCACAAGAATCATAACACGA<br>GCCACAAGAAGCCGATGCAAGCGGAGCCGGACCGCGGCACCAAGGCGCGGAACACGA<br>CGCCAGGCCAAGCAGAAGCTCGCCGGCGGCCAGCAAGGTCCACAGCGGCGCGGCCAC<br>CAGTGCACTCTATGTTGAGAGTGGACCAACGTTGACGCCGAAGGCTCACAATGGCTGAG<br>CGCTGTAAGTGGCGTCGCGCGCCAGGTTGCCACACCAGGAACAGACGCAACAACCACG<br>AGCGGCCGTACAAGCATAGAGCCGTGCCTGAATGGTTGATCCAGTAGACGGCCACAAG<br>GCAGGTGGCGGTCGAACCGAATGGCTAGGTCCGGCTCACAAGGATAACACGGCTACAC<br>CTGAACGCCGCCGCAGGCGCAACGACGAAGCCGTAGCAACGCAGAGAGATAACGCAAC<br>GGCCAGGCAAGAGATACACAATTTCATGTCTTGACGGTCCTCGTGAACTGTG**GTTAAC** |

**Supplementary Table 20**. Template sequences used in this study. Underlined sequence denotes T7 promoter; Underlined bold sequence denotes restriction enzyme sites used for template linearization (HpaI for RNA1, RNA5, RNA7, RNA8 and

RNA9; NotI for RNA2; NheI for RNA6). For SP6 promoter sequence, the T7 promoter sequence was replaced with ATTTAGGTGACACTATA.

**Supplementary Figure 1**

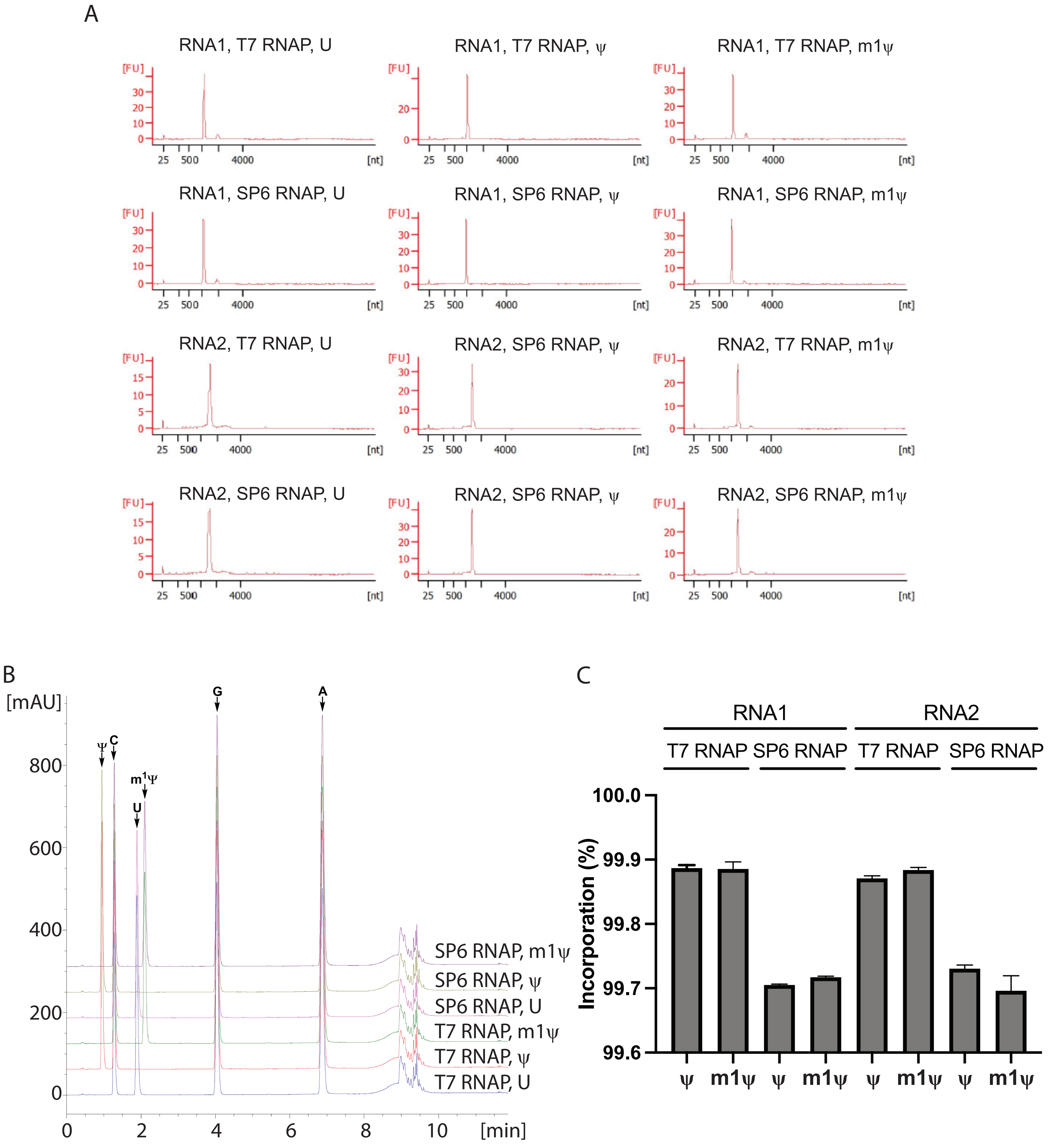

**Supplementary Figure 2**

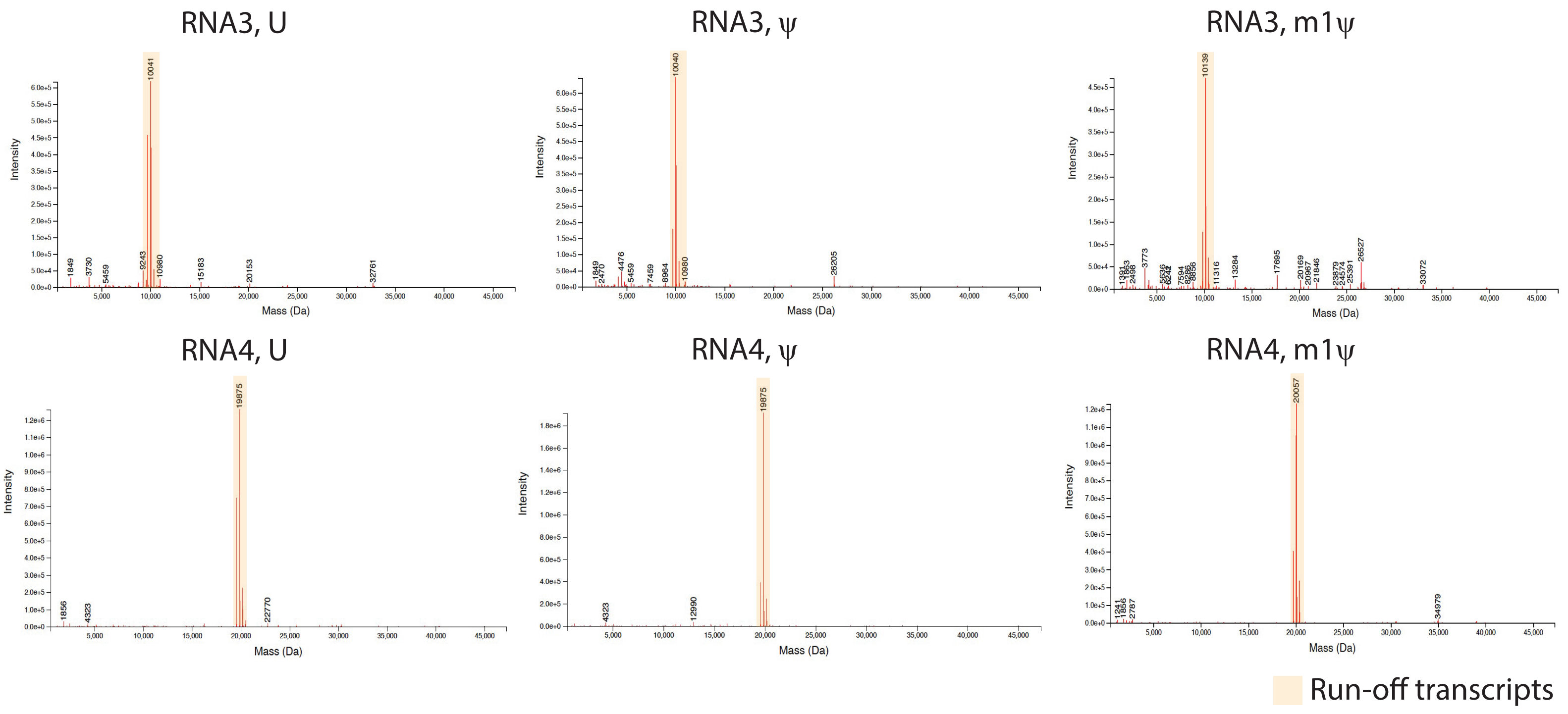

**Supplemental Figure 3.**

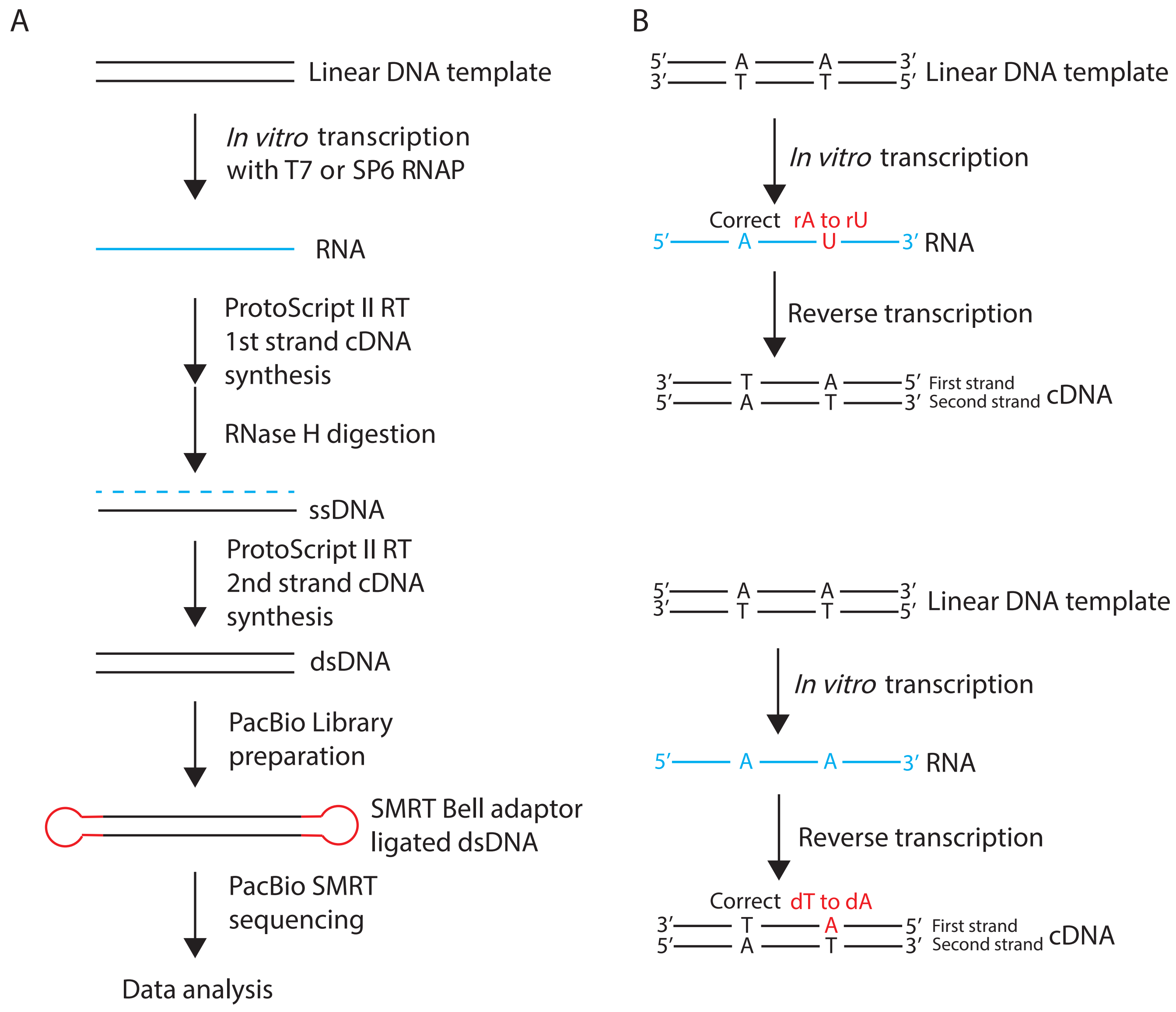

**Supplemental Figure 4**

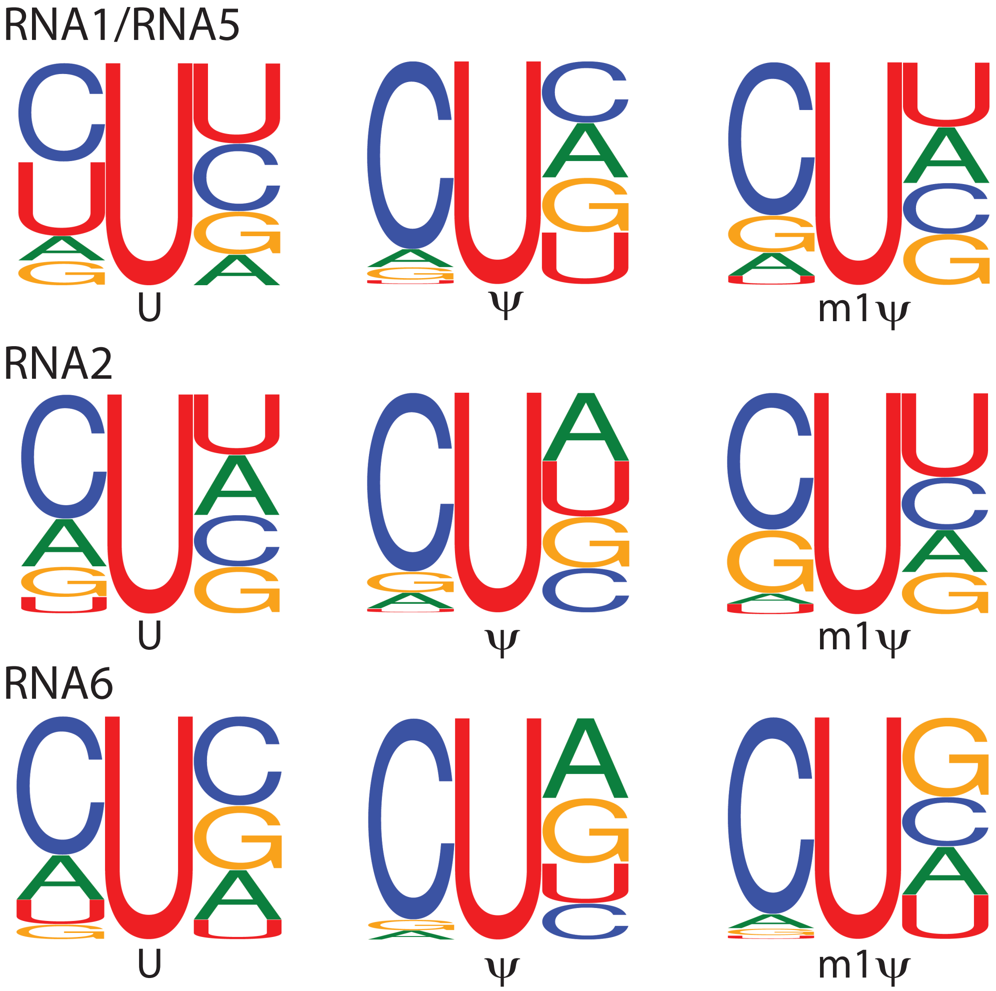

**Supplemental Figure 5**

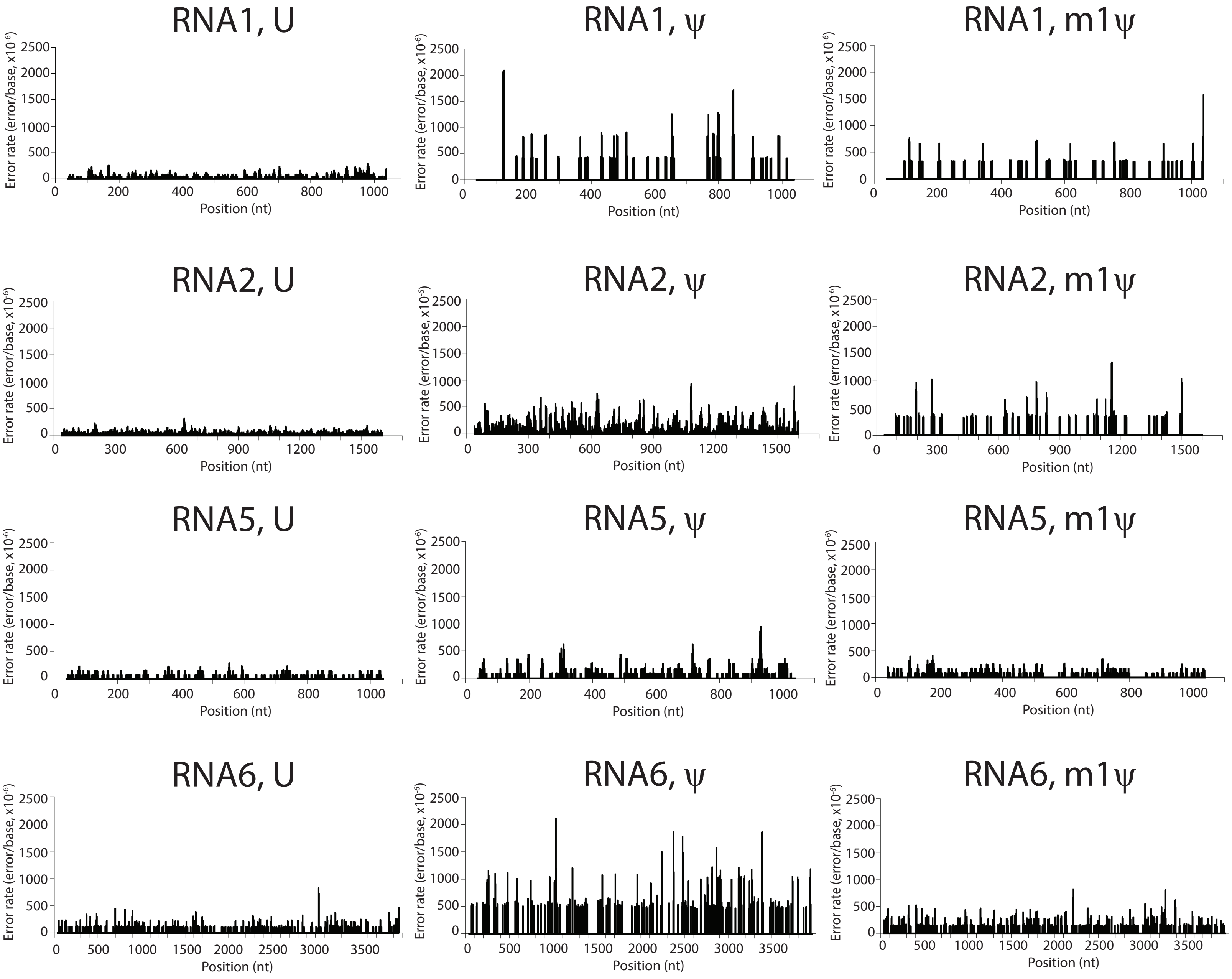

**Supplemental Figure 6**

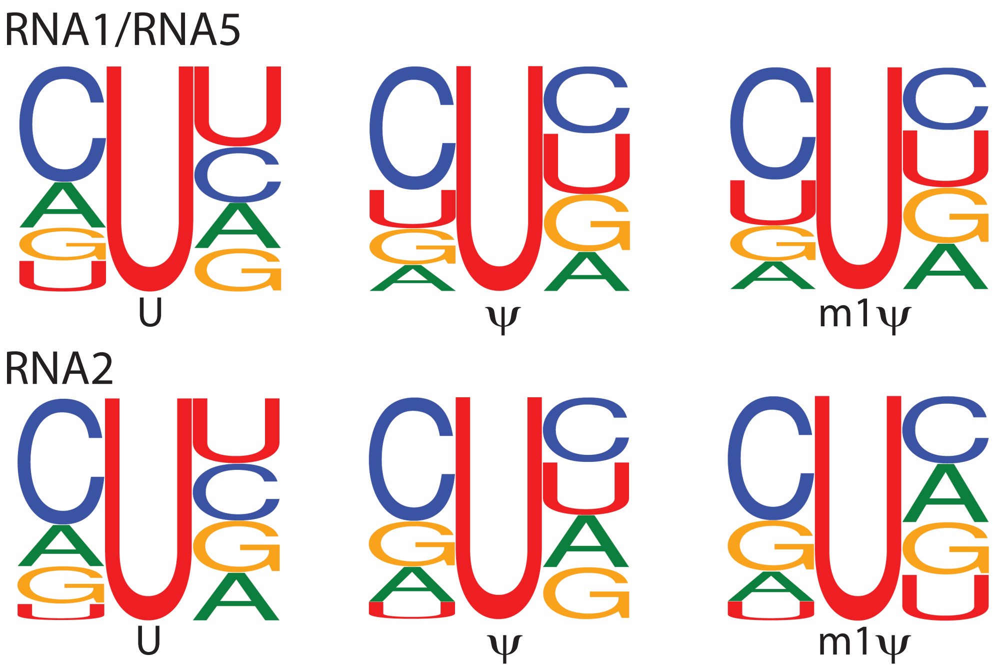

**Supplemental Figure 7**

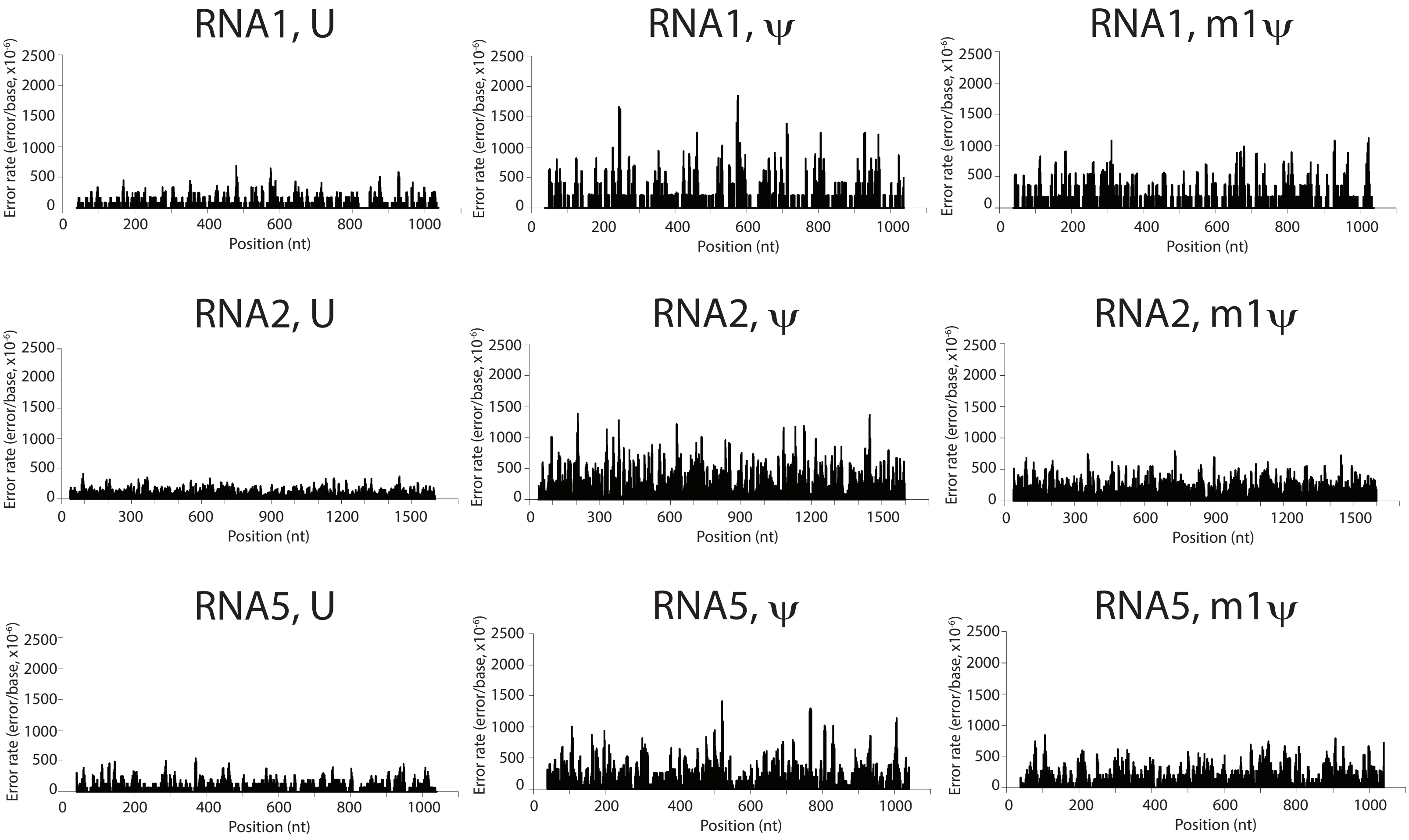

### Supplemental Figure 8

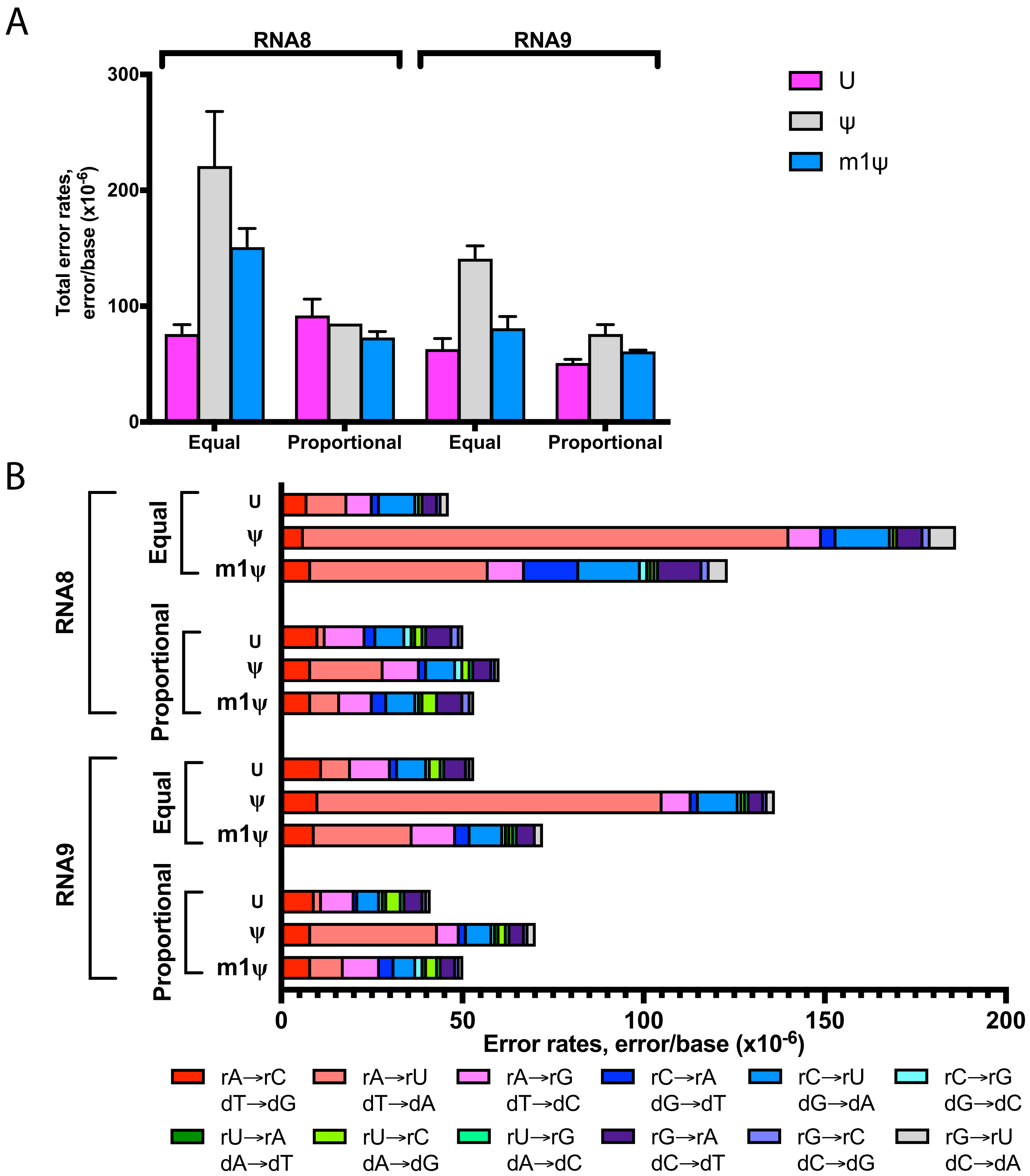

### Supplemental Figure 9

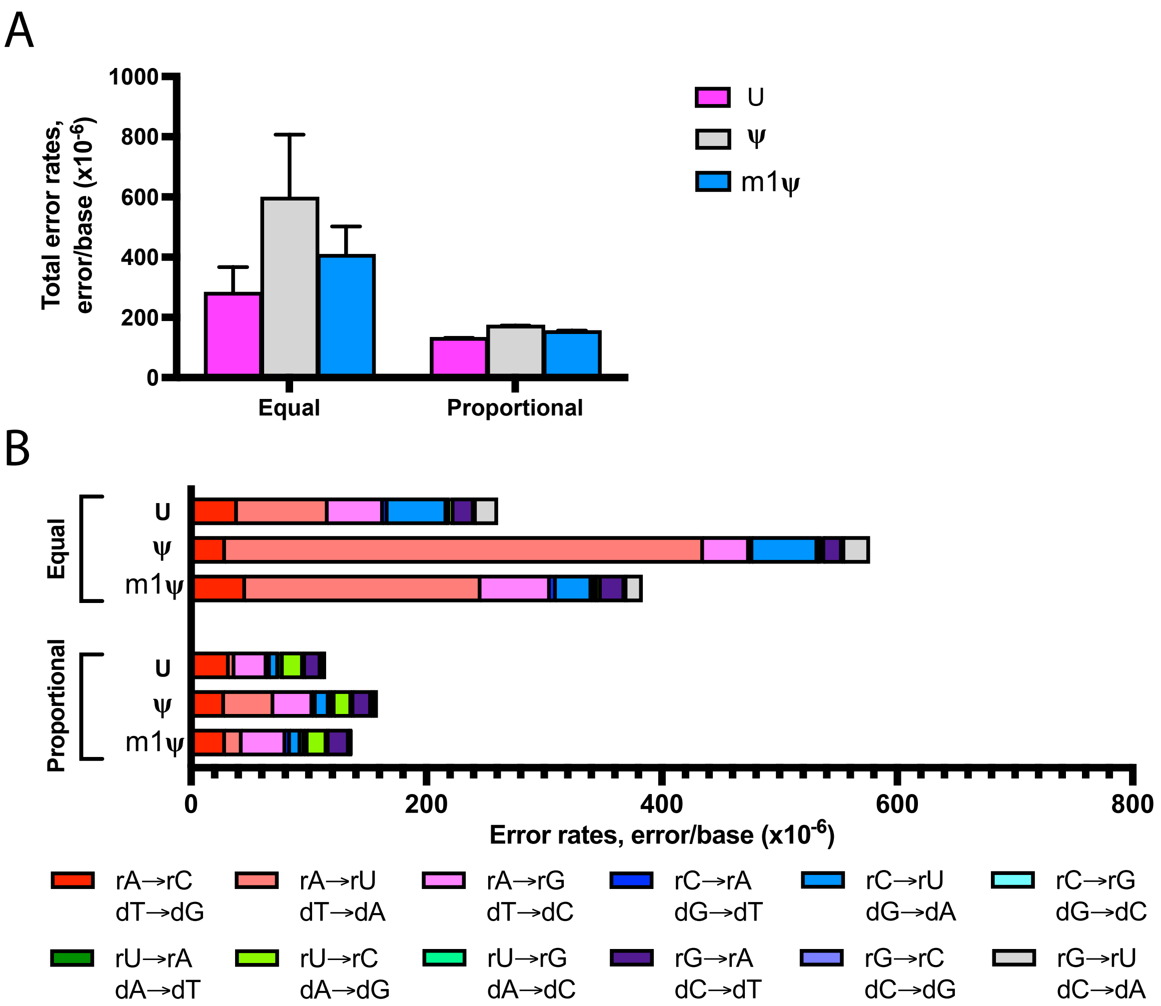

### Supplemental Figure 10

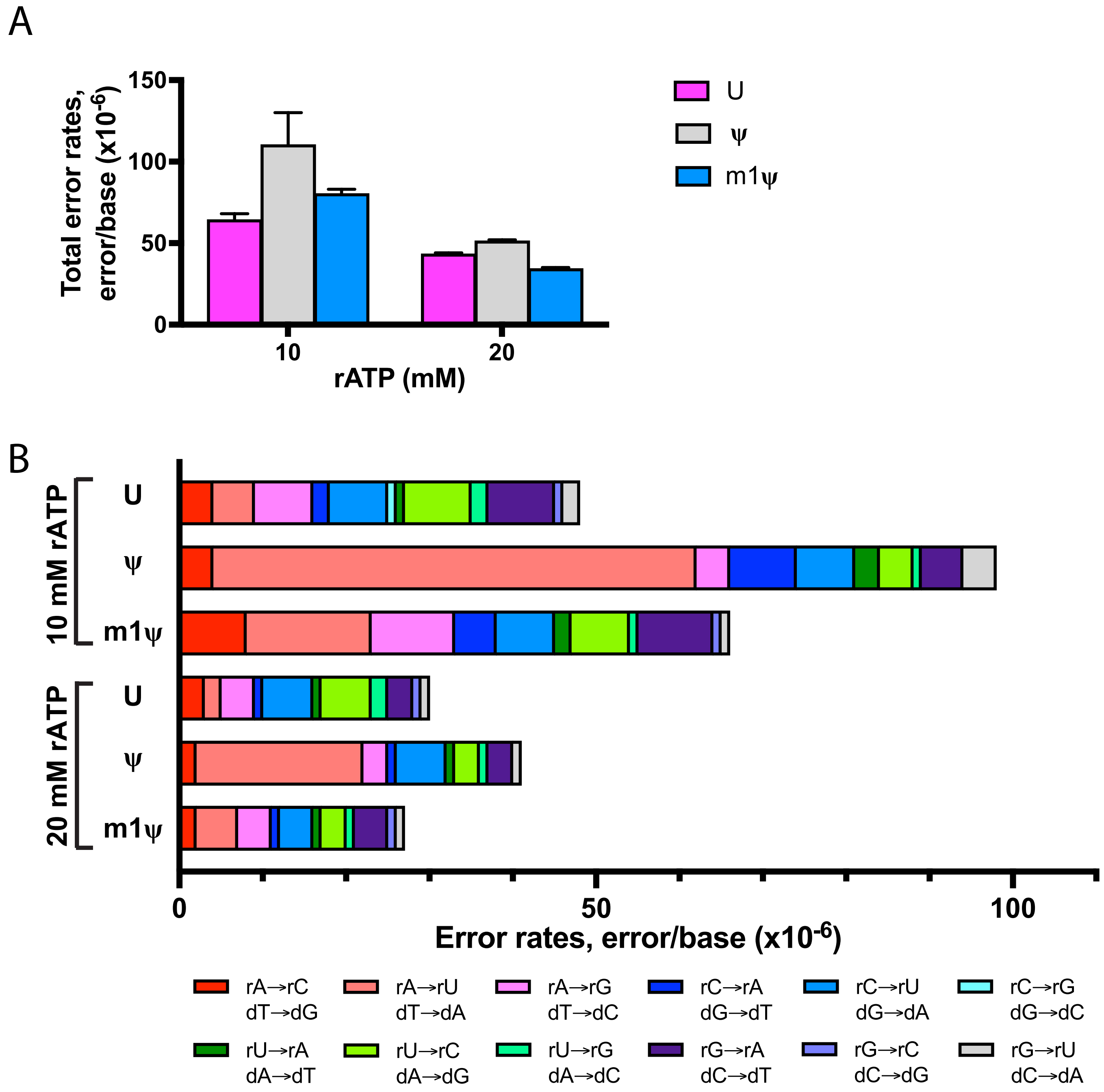
